# A *Vibrio cholerae* Core Genome Multilocus Sequence Typing Scheme to Facilitate the Epidemiological Study of Cholera

**DOI:** 10.1101/2020.01.27.919118

**Authors:** Kevin Y. H. Liang, Fabini D. Orata, Mohammad Tarequl Islam, Tania Nasreen, Munirul Alam, Cheryl L. Tarr, Yann F. Boucher

## Abstract

Core genome multilocus sequence typing (cgMLST) has gained popularity in recent years in epidemiological research and subspecies level classification. cgMLST retains the intuitive nature of traditional MLST but offers much greater resolution by utilizing significantly larger portions of the genome. Here, we introduce a cgMLST scheme for *Vibrio cholerae*, a bacterium abundant in marine and freshwater environments and the etiologic agent of cholera. A set of 2,443 core genes ubiquitous in *V. cholerae* were used to analyze a comprehensive dataset of 1,262 clinical and environmental strains collected from 52 countries, including 65 newly sequenced genomes in this study. We established a sublineage threshold based on 133 allelic differences that creates clusters nearly identical to traditional MLST types, providing backwards compatibility to new cgMLST classifications. We also defined an outbreak threshold based on seven allelic differences that is capable of identifying strains from the same outbreak and closely related isolates which could give clues on outbreak origin. Using cgMLST, we confirmed the South Asian origin of modern epidemics and identified clustering affinity among sublineages of environmental isolates from the same geographic origin. Advantages of this method are highlighted by direct comparison with existing classification methods, such as MLST and single nucleotide polymorphism-based methods. cgMLST outperforms all existing methods in terms of resolution, standardization, and ease-of-use. We anticipate this scheme will serve as a basis for a universally applicable and standardized classification system for *V. cholerae* research and epidemiological surveillance in the future. This cgMLST scheme is publicly available on PubMLST (https://pubmlst.org/vcholerae/).

**IMPORTANCE:** Toxigenic *Vibrio cholerae* of the O1 and O139 serogroups are the causative agent of cholera, an acute diarrheal disease that plagued the world for centuries, if not millennia. Here, we introduce a core genome multilocus sequence typing (cgMLST) scheme for *V. cholerae*. Using cgMLST, we established an outbreak threshold that can efficiently identify outbreak related strains and potential sources of introduction. We also defined a sublineage threshold that is similar to traditional MLST sequence type which will provide context to this new typing method by relating it to previous MLST results. cgMLST outperforms all existing methods in terms of resolution, standardization, and ease-of-use, making this scheme the most suitable method for *V. cholerae* typing and surveillance worldwide.

## INTRODUCTION

Cholera is transmitted in a fecal-oral route mostly by contaminated food or water (1, 2). The case fatality rate (CFR) of this disease can be up to 50% without treatment, but with proper medical care, the CFR is usually less than 1% (2, 3). In developed countries with proper water treatment facilities, cholera is practically non-existent aside from imported cases. Unfortunately, this cannot be said for many developing countries lacking this infrastructure, where cholera has been endemic for centuries such as in parts of South Asia (4). As it is also difficult to eradicate cholera (5), this disease often becomes endemic in regions where it has been introduced, for example in Latin America in 1991 (6, 7), Haiti in 2010 (8), and Yemen in 2016 (9). It is estimated that there are over a million cholera cases each year resulting in tens of thousands of deaths worldwide (10). Being an indicator of healthcare and socio-economic disparities (11, 12), this disease is often under-reported due to its negative influence on tourism as it implies poor water quality (13). Together with the lack of a universally applicable and standardized classification method, outbreak surveillance and source attribution is often challenging (1, 8). The Haiti outbreak for example, due to these limitations, required extensive genomic and epidemiological research since the beginning of the outbreak to determine the source of introduction, which was not confirmed until August 2011 even though cholera broke out in July 2010 (8, 14–17).

A typing method for use in global surveillance of pandemic causing pathogens such as *V. cholerae* should be efficient and easy to use, with the potential to be applied to all *V. cholerae* strains around the world. Therefore, it must have the capacity to analyze thousands of genomes efficiently and new genomes should be easily typed as they get sequenced. As all cholera outbreaks are caused by a single lineage of *V. cholerae*, the pandemic generating/phylocore genome (PG) lineage, which includes the 7^th^ pandemic El Tor, El Tor sister, El Tor progenitor, Classical, and Classical sister clades (5, 18, 19), this method should also be able to differentiate isolates at a fine scale and separate outbreaks caused by genetically similar strains. Such a method will help create a comprehensive database with detailed epidemiological data that will allow for the analysis of future outbreak strains in a global context and guide subsequent epidemiological analyses. Different methods for subspecies level classification and outbreak surveillance have been developed for *V. cholerae*. These methods include serotyping, multilocus sequence typing (MLST) (20, 21), multilocus variable number of tandem repeats (VNTR) analysis (MLVA) (22, 23), and single nucleotide polymorphism (SNP)-based approaches (14). Despite the popularity of these methods, there are important limitations to each.

Serotyping based on the presence of cell surface O-antigens is one of the earliest attempts at subspecies level classification of *V. cholerae*. There are now over 200 serogroups of *V. cholerae* identified; however, only the toxigenic members of the O1 and O139 serogroups have been found to be responsible for all major documented epidemics and pandemics (24, 25). Serogroup O1 can be further divided into two biotypes (El Tor and Classical) and three serotypes (Inaba, Hikojima, and Ogawa) (2). The lack of resolution within the epidemic strains and the possibility of serogroup conversion (26) limits the use of serotyping in epidemiological studies.

MLST provides a standardized classification method that is based on a collection of six to seven well-defined housekeeping genes (27). MLST was used to study a number of cholera outbreaks and allowed the descriptions of its general population structure (28, 29). It is reproducible and provides reliable results; however, it is unable to differentiate between closely related strains which limits its use in outbreak surveillance (30, 31). In addition, direct comparisons between different MLST schemes are difficult, as different schemes utilize different housekeeping genes.

MLVA utilizes VNTR regions, which are under less selective pressure than housekeeping genes. This method therefore provides greater resolution than MLST for some bacterial species (32, 33). However, due to their rapid mutation rate, VNTR regions are more affected by homoplasy where two isolates may share the same MLVA profile due to convergent mutation and not by vertical descent (34). As a result, MLVA may produce clusters that do not necessarily reflect phylogenetic relationships (35). Two common PCR-based methods exist for the typing of VNTR regions, but each have significant limitations (36). The first method is multiplex PCR which can analyze all loci at once, but it is impossible to determine which bands correspond to which loci; therefore, this method only produces a banding pattern for strain identification, which makes it difficult to standardize and communicate results. The second method is the separate amplification of VNTR regions but determining the number of repeats based on amplicon size information alone is difficult if the difference in size is not large enough. In addition, different types of mutations that do not necessarily change the number of repeats can cause a change in amplicon size. Sequencing is needed to confirm MLVA profiles, but repeat regions increase the chances of sequencing errors (37). Due to these limitations, stringent quality control is required for reliable MLVA analysis (38).

SNP-based analysis is one of the most common whole genome-based methods currently being used and was applied to various outbreaks (14, 39, 40). It relies on the identification of conserved SNPs in strains of interest using next-generation sequence reads or assembled genomes. The number of SNPs can then be related to the evolutionary distance between isolates. SNP-based analysis provides reliable results with sufficient resolution for epidemiological studies, but it is sensitive to horizontal gene transfer and recombination events, as each event will result in many SNPs being created. The number of SNPs between two strains, therefore, does not necessarily reflect true phylogenetic relationship. SNPs found in recombinogenic regions should therefore be removed which, depending on the organism of interest, can be anywhere from 30% to 97% of SNPs identified (41, 42). Since recombination and horizontal gene transfer events are common within *V. cholerae* (43–45) and between the species and its close relatives (46, 47), SNP-based methods, although suitable in individual epidemiological studies, will be difficult to serve as a universal classification method for *V. cholerae*.

Core genome MLST (cgMLST), also known as the gene-by-gene approach, overcomes the various limitations of the previously mentioned subtyping methods and was established to serve as a universally applicable standardized typing scheme. Similar to MLST, cgMLST relies on individual gene sequences to differentiate between closely related strains; however, instead of using only six to seven housekeeping genes, cgMLST utilizes hundreds to thousands of core genes, which are commonly found in all strains of a species. By utilizing a much larger portion of the genome, cgMLST provides superior resolution compared to traditional MLST schemes. By combining the expandable and standardized classification method that made traditional MLST favourable with the resolution of whole genome-based methods, cgMLST is becoming more popular in epidemiological and ecological studies (48–54). This method has the added advantage of backwards compatibility with all MLST schemes. This means that it is possible to determine MLST profiles of any isolates based on their cgMLST profiles, since cgMLST would include all housekeeping genes by definition. This allows for a 1:1 mapping of any previously established MLST scheme to the cgMLST scheme, helping consolidate already existing classification information.

Another major benefit of cgMLST is that, much like traditional MLST methods, it is possible to establish different clustering thresholds to define important groups. Clonal complexes are examples of clustering thresholds established by MLST schemes, where each clonal complex corresponds to a cluster of isolates that share at most one allelic difference across all seven genes sequenced. Some important clonal complexes were shown to correspond to either groups established by a previous typing method (55) or major outbreak strains (56). However, cgMLST offers even greater flexibility than MLST in this regard given the number of loci considered. With large clustering thresholds, it is possible to identify lineage- or even sublineage-level differences to study large scale patterns and answer broader ecological questions. Furthermore, with smaller clustering thresholds where groups are created based on the sharing of a larger number of alleles, it is possible to identify very closely related strains useful in epidemiological studies. The benefits of defining clustering thresholds through cgMLST have already been demonstrated in other human pathogens, such as *Brucella melitensis* (52), *Campylobacter jejuni* (51), *Clostridium difficile* (53), *Enterococcus faecium* (50), and *Listeria monocytogenes* (49).

In this study, we introduce a cgMLST scheme for the genome-wide typing of *V. cholerae* and demonstrate its universality and efficacy by applying it to known cholera outbreaks around the world. The advantages of cgMLST are presented by comparing the scheme with previously established classification methods. Additionally, we have produced a 1:1 mapping of the cgMLST scheme against two previously established MLST schemes for *V. cholerae* (20, 21), allowing for the consolidation of existing classification information. The cgMLST scheme, genome sequences used in this study, and relevant epidemiological information are publicly available on PubMLST (https://pubmlst.org/vcholerae/), which allows for the automatic annotation and subsequent analyses of hundreds of newly uploaded *V. cholerae* genomes in a global context. This increase in efficiency, standardizability, and resolution compared to current methods make cgMLST the most suitable classification scheme for large scale *V. cholerae* surveillance. By applying this scheme to our collection of over 1,200 isolates collected around the world, it was possible to establish outbreak and sublineage thresholds which not only allowed us to validate the South Asian origin of many modern epidemics as proposed in previous studies (5, 57, 58) but also identify clustering affinity among environmental strains, where isolates from the same sublineage are also likely from the same geographic region. This pattern is not seen in clinical isolates, as human hosts readily carry them over large geographical distances.

## RESULTS AND DISCUSSION

### A high-resolution typing scheme for pandemic *V. cholerae*

The highest level of resolution of any cgMLST scheme is defined by core genome sequence types (cgSTs), where a unique cgST represents a unique allelic profile. Isolates that belong to the same cgST are expected to be phylogenetically very closely related, as although they may not have the exact genomic sequence, they do have the same sequence for all 2,443 core gene loci used in cgMLST. We identified a total of 1,026 cgSTs from 1,262 genomes collected from 52 countries. Even with our extensive dataset, we have yet to sample anywhere close to the total predicted cgST diversity for the global *V. cholerae* population (Fig. S1). All isolates were given at least one cgST designation and up to two MLST sequence type (ST) designations based on two previously established MLST schemes (20, 21) (Table S1). MLST STs are defined based on the unique combination of all loci of a particular MLST scheme, which ideally uses six to seven well-defined housekeeping genes. Only 12 STs are exclusively present in the 7^th^ pandemic El Tor lineage identified using traditional MLST (20, 21), whereas 560 cgSTs are uniquely present in this group based on cgMLST (Table S1). As the El Tor lineage is responsible for most cholera outbreaks around the world since the beginning of the 7^th^ pandemic (59), this superior ability to resolve between closely related strains in the 7^th^ pandemic El Tor lineage makes cgMLST more suitable in outbreak surveillance than traditional MLST.

### Backwards compatibility with previous subspecies classification methods

Much like how cgSTs are important in studying closely related strains typical in outbreaks, it is also important in establishing a standardized nomenclature at a higher level to answer broader ecological questions. Here, we propose a sublineage definition for *V. cholerae* based on cgMLST.

Pairwise allelic differences calculated between all isolates show three major peaks (Fig. 1A). The first peak ends at 40 allelic differences, and the second peak ends at 133 allelic differences (Fig. 1B). The last peak begins at 2,200 allelic differences (Fig. 1A), which is expected due to mutational saturation (i.e., every single allele in the scheme is different between the two distantly related strains being compared). Both breaks (i.e., 40 or 133 allelic differences) could represent a potential sublineage delineation. To choose between the two thresholds, the clustering efficiency is measured by calculating the Dunn Index (DI) (60) (see Materials and Methods). Since cluster distances are measured by allelic differences, the network with the best clustering efficiency (i.e., the highest DI) will also produce clusters that best represent biological relationships, as isolates are more closely related to themselves than to isolates from other clusters. A DI was calculated for each clustering threshold in the range of 1 to 1,000 allelic differences with 100 bootstrap replicates (Fig. 2). As the clustering threshold defines the maximum number of allelic differences within a cluster, the smaller the threshold, the more closely related the isolates are within a cluster. It is clear that DIs in the range of 0 to 50 allelic differences are significantly lower than the DIs in the range of 100 to 350 allelic differences, with 133 being a clear local maximum. Since 133 allelic differences has the best clustering efficiency and also represents a natural break where most isolate pairs have either less than or much greater number of allelic differences (Fig. 1B), it was therefore chosen as the sublineage threshold.

**Fig. 1.**
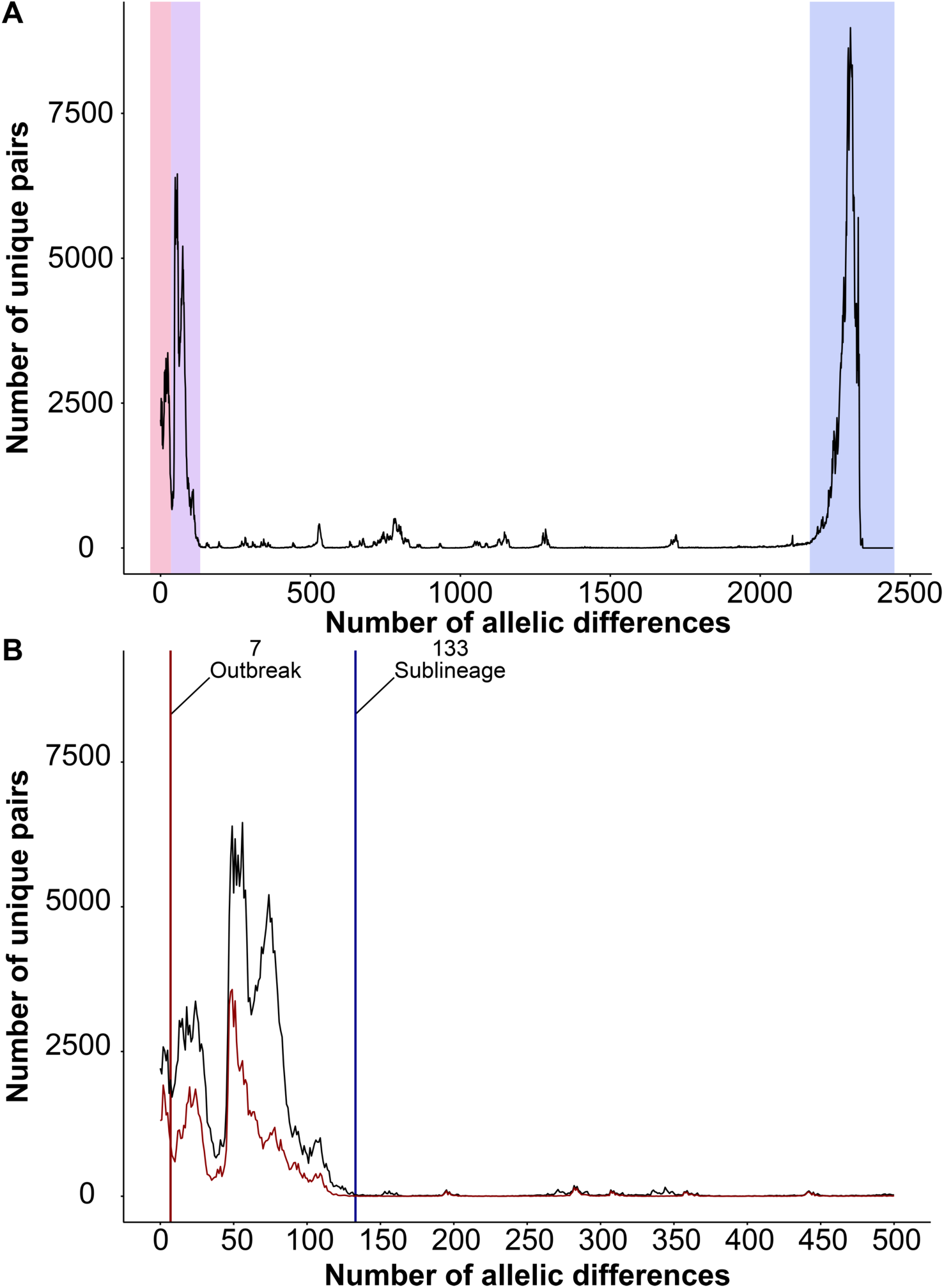
Pairwise allelic differences for all isolates used in this study. Both plots show the frequency of allelic mismatches in pairwise comparisons. A) Pairwise comparisons of up to 2,443 allelic differences are shown. Major peaks are shaded. B) Comparisons with up to 500 allelic differences are shown. Pairwise comparisons of only clinical isolates are shown in red. Vertical lines indicate the outbreak threshold (red) and sublineage threshold (blue).

**Fig. 2.**
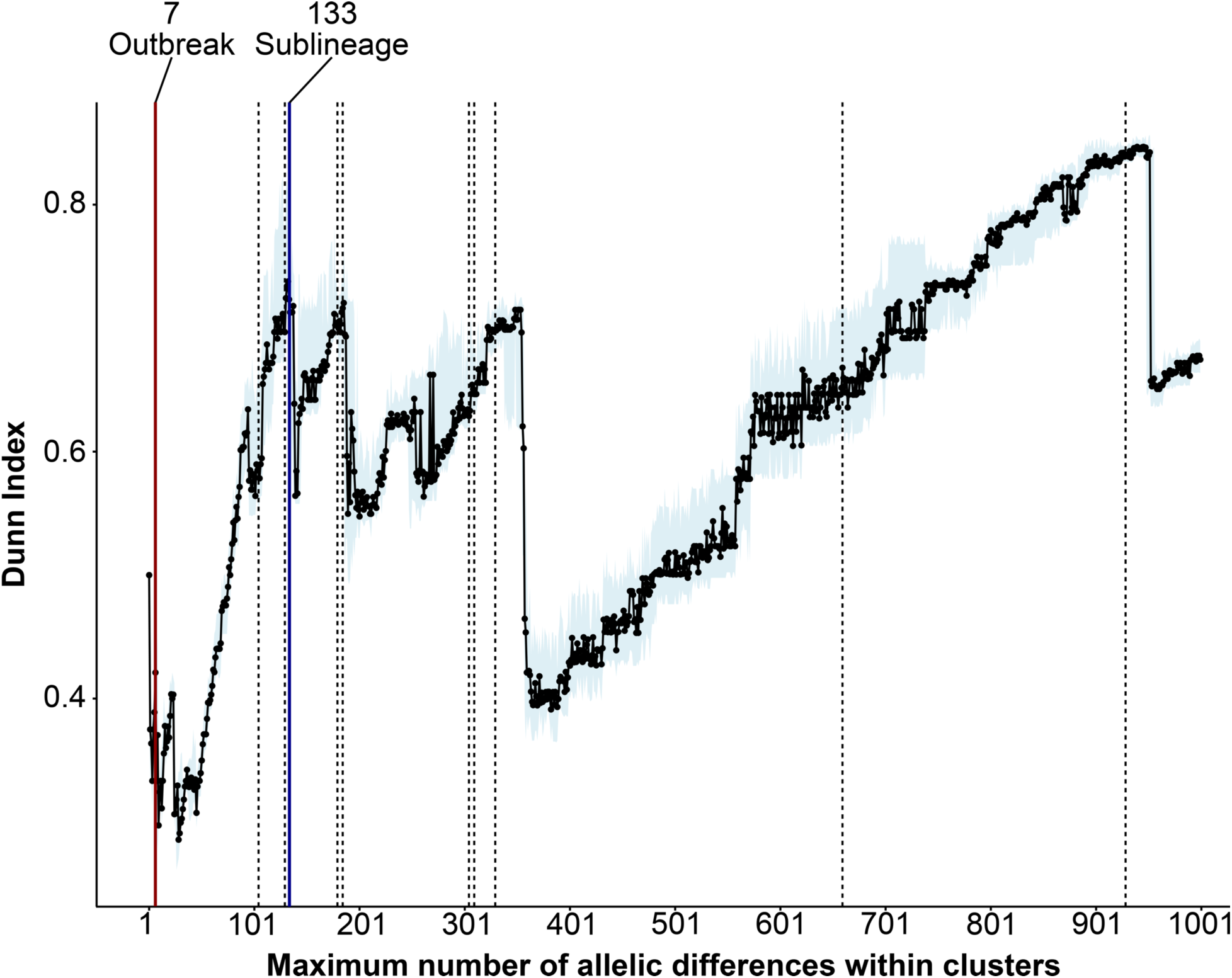
Plot showing the Dunn Index for clustering thresholds ranging from 1 to 1,000 allelic differences. Each clustering threshold is bootstrapped 100 times. The median, plotted with the light blue shade, indicates the 25^th^ to 75^th^ percentile range. Red and blue vertical lines indicate the outbreak and sublineage thresholds, respectively. The dotted lines represent other clustering thresholds used in the Adjusted Rand Index calculations (Fig. 3B and Fig. S3).

Because cgMLST includes all housekeeping genes, information from the two MLST schemes previously developed for *V. cholerae* (20, 21) can now be consolidated with the cgMLST scheme by creating a 1:1 cgMLST to MLST map. To evaluate the similarities between the sublineage threshold and the MLST schemes, we created a minimum spanning tree (MST) for all Bangladesh isolates (*n* = 255) showing only edges with 133 allelic differences or fewer (Fig. 3A and Fig. S2). Each cluster therefore represents a single sublineage. Bangladesh was chosen to compare cgMLST and MLST as it is the most extensively sampled country both in terms of clinical and environmental isolates in our dataset. Using this dataset, the chosen sublineage threshold produces clusters that closely resemble ST clustering from traditional MLST. Based on the 2013 MLST scheme (20), each sublineage corresponds to exactly one ST (Fig. S2), whereas there is only one sublineage that contains two STs based on the 2016 MLST scheme (21) (Fig. 3A). All but two isolates belong to ST1; N16961 and A19 belong to ST290, which differs from ST1 at only one of seven MLST loci (Table S2). The reason these two isolates are of a different MLST ST could only be partly explained; they were isolated at an earlier time point (1970s near the start of the 7^th^ pandemic (61)) than most of the remaining isolates, which were isolated from 1991 onwards (Table S1).

**Fig. 3.**
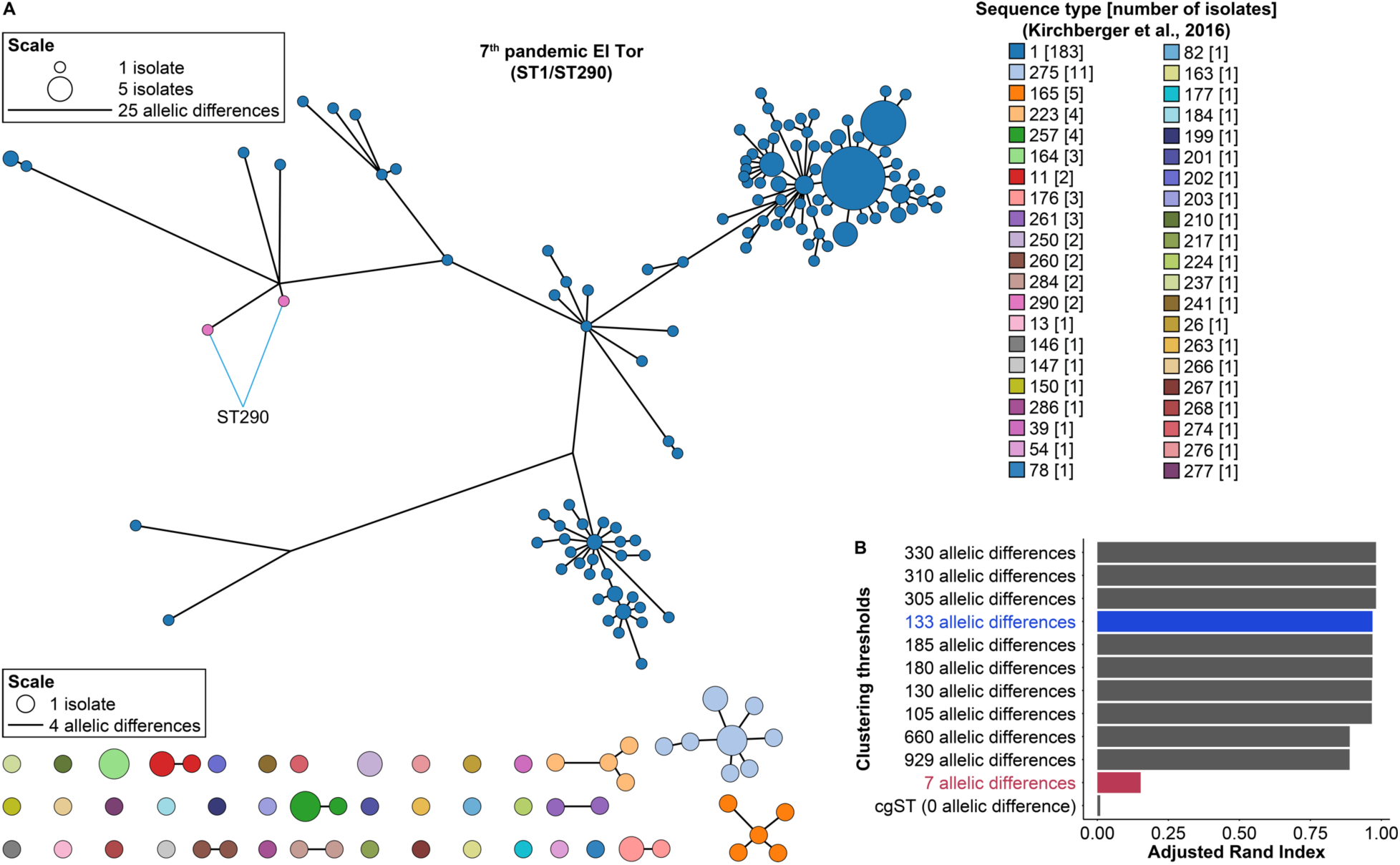
Evaluation of network similarities between the cgMLST sublineage threshold and MLST ST. A) Networks of all sublineages identified using only *V. cholerae* isolates from Bangladesh (*n* = 255). Each cluster represents a sublineage and includes isolates with less than or equal to 133 allelic differences with each other. Each node represents a cgST and is colored by sequence type based on the 2016 MLST scheme (21). Size of the nodes are proportional to the number of isolates. The length of the connecting lines within a cluster is proportional to the number of allelic differences. B) Adjusted Rand Index for individual pairwise comparisons between predefined clustering thresholds (Fig. 2) and the 2016 MLST scheme (21). The sublineage clustering threshold (i.e., 133 allelic differences) and outbreak threshold (i.e., 7 allelic differences) are indicated in blue and red bars, respectively.

It is impossible to visually evaluate similarities between two MSTs with over 1,200 nodes each simply due to the sheer volume of data. The Adjusted Rand Index (ARI) was therefore used as a metric to determine network similarities (62) (see Materials and Methods). In order to determine whether the sublineage threshold (i.e., 133 allelic differences) is indeed the best match to traditional MLST schemes, we chose 11 clustering thresholds distributed across the range of 1 to 1,000 allelic differences (Fig. 2) to compare with the MLST schemes. These additional thresholds are chosen as they have a relatively high DI compared to their immediate neighbours. More data points were chosen in the range of 105 to 330 allelic differences, as it was expected that thresholds in this range will best match the traditional MLST schemes. Interestingly, all thresholds in that range had comparable ARIs when compared to both the 2016 (21) and the 2013 MLST schemes (20) (Fig. 3B and Fig. S3), indicating that all of them, including the sublineage threshold, produce clusters similar to the MLST scheme. This would suggest that there can be a large range of diversity within a single MLST ST where isolates can have anywhere from 0 (i.e., have the same cgST) to 330 allelic differences. Although clustering thresholds between 105 to 330 allelic differences produce similar clusters to a traditional MLST scheme, the sublineage threshold of 133 allelic differences was chosen as it has the best clustering efficiency (Fig. 2) and it represents a natural breakpoint in the currently sampled population (Fig. 1B).

A phylogenetic tree of 1,146 isolates was used to assess the phylogenetic support of the sublineage threshold across different *V. cholerae* strains (Fig. 4). This tree includes all *V. cholerae* isolates within our dataset with the exception of the recently published 116 clinical isolates from the Yemen cholera outbreak (9), which all belongs to the 7^th^ pandemic El Tor lineage. The strains within the PG lineage are closely related with little genetic variation. These lineages are therefore collapsed in the phylogenetic tree as the relationships between them are not well resolved. All sublineages formed monophyletic clades, although in some cases the most basal branch is of a different sublineage (e.g., *V. cholerae* strains T5 or 506315) creating paraphyletic clades. Ideally, each sublineage would correspond to exactly one full monophyletic clade. The reason this is not seen is likely the lack of sampling, leading to the grouping of relatively distantly related isolates together in the same clade. Further sampling will likely resolve these cases into two separate monophyletic clades. Out of 1,262 isolates, we identified 291 sublineages, 19 of which belong exclusively to the PG lineage and 223 are singletons. Much like cgSTs, the rarefaction curve also indicates that the total sublineage diversity of *V. cholerae* is far from being sampled (Fig. S1).

**Fig. 4.**
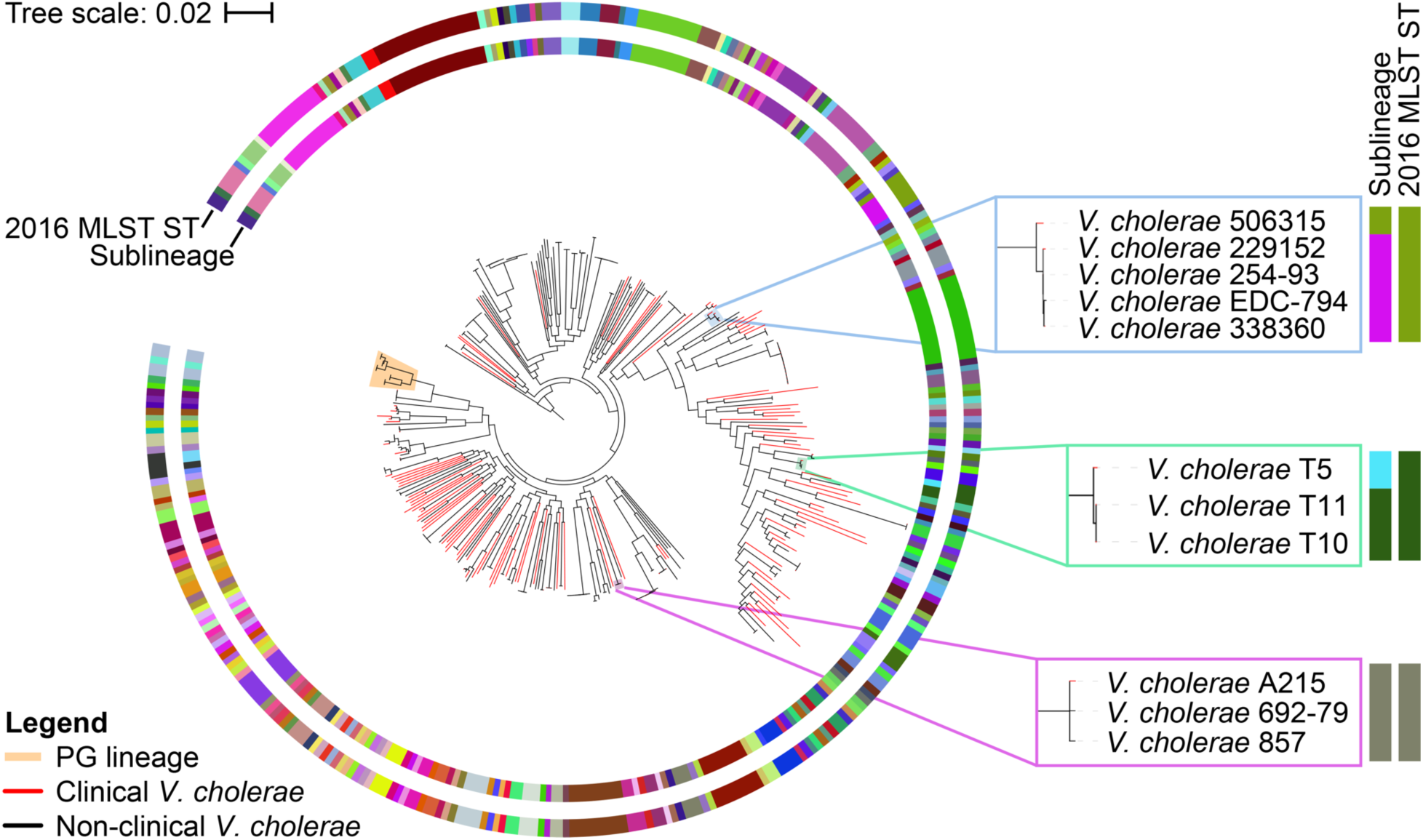
Phylogenetic tree of 1,146 *V. cholerae* isolates (excluding the 116 isolates from the Yemen cholera outbreak) reconstructed using Parsnp v1.2 (92). All group inside the PG lineage (7^th^ pandemic El Tor, El Tor progenitor, El Tor sister, Classical, and Classical sister) are collapsed. Outer rings represents clustering by sequence type based on the 2016 MLST scheme by Kirchberger and colleagues (21), whereas the inner ring represents clustering based on the sublineage threshold (i.e., 133 allelic differences). Branches of clinical strains are colored in red. The phylogenetic tree is rooted with a basal lineage to *V. cholerae* (collapsed) (77, 78).

The sublineage concept has been applied to numerous pathogens and as such were each defined differently depending on the pathogen in question. Some have defined sublineages based on natural breaks in genetic similarities (49), while others may use sublineage to refer specifically to traditional MLST STs (63) or even finer level of resolution below the MLST ST level based on whole genome analyses (64). There is, however, one unifying feature of all sublineage definitions – that they all refer to monophyletic clades. Sublineages are defined in this study based on natural breaks in allelic differences calculated from cgMLST profiles and were put into context by comparing with two traditional MLST schemes. We have shown that our definition of sublineage results in monophyletic clades but also corresponds to any traditional MLST ST designation (Fig. 4). This sublineage definition will therefore play a crucial role in consolidating information from all previous MLST analyses.

### A universal South Asian origin for modern cholera outbreaks

With the continual improvement of next-generation sequencing techniques, whole genome sequencing is expected to become a standard practice or even the first identification tool used in clinical and epidemiological studies. It is therefore critical to develop a rapid typing scheme for genome sequence data that has the power to inform us about the relationship of a novel isolate with known strains. This is done here by defining what we term an ‘outbreak threshold’ based on cgMLST, which can identify outbreak related strains and potential sources of introduction. The outbreak threshold is expected to be less than 40 allelic differences as isolates from the same outbreak are very closely related (9, 14). There is a minor discontinuity at seven allelic differences where most isolate pairs have either less or more than this number of allelic differences (Fig. 1B). Looking at the DI, the local maximum in the range of 0 to 50 occurs at seven allelic differences as well (Fig. 2), making this cutoff a likely candidate for an outbreak threshold. When applying the outbreak threshold to a full dataset containing all sequenced *V. cholerae* genomes meeting the minimum quality threshold, major clusters were examined to evaluate the ability of cgMLST to identify strains that are part of the same outbreak.

One of the major outbreak clusters identified, with no prior information required, contains the Haiti and the Yemen outbreaks, which are the two best documented cholera outbreaks in modern history (8, 9, 14, 16, 65). Isolates collected from these outbreaks form a single cluster with the Dominican Republic, Eurasian (India, Russia, Nepal, and Ukraine), and African (Tanzania, Kenya, and Somalia) isolates (Fig. 5A). The Dominican Republic isolates are closely related to the Haiti outbreak strains. Given the close proximity of the two countries, co-located on the island of Hispaniola, it was expected that isolates from Haiti would eventually spread to the Dominican Republic (14). The 7^th^ pandemic El Tor lineage spread across the world from South Asia in three separate waves (61). The third wave, being the most recent distribution event, has been claimed to be responsible for the outbreaks in Haiti and Yemen (9). It is therefore not surprising to see Haiti and Yemen form a single cluster with India (i.e., South Asia) at its center. Nepal is the known source of introduction for the Haiti outbreak in 2010 (16), and comparison with over 1,200 *V. cholerae* isolates from all over the world still shows the Nepalese isolates as the closest relatives to the Haitian isolates (Fig. 5A).

**Fig. 5.**
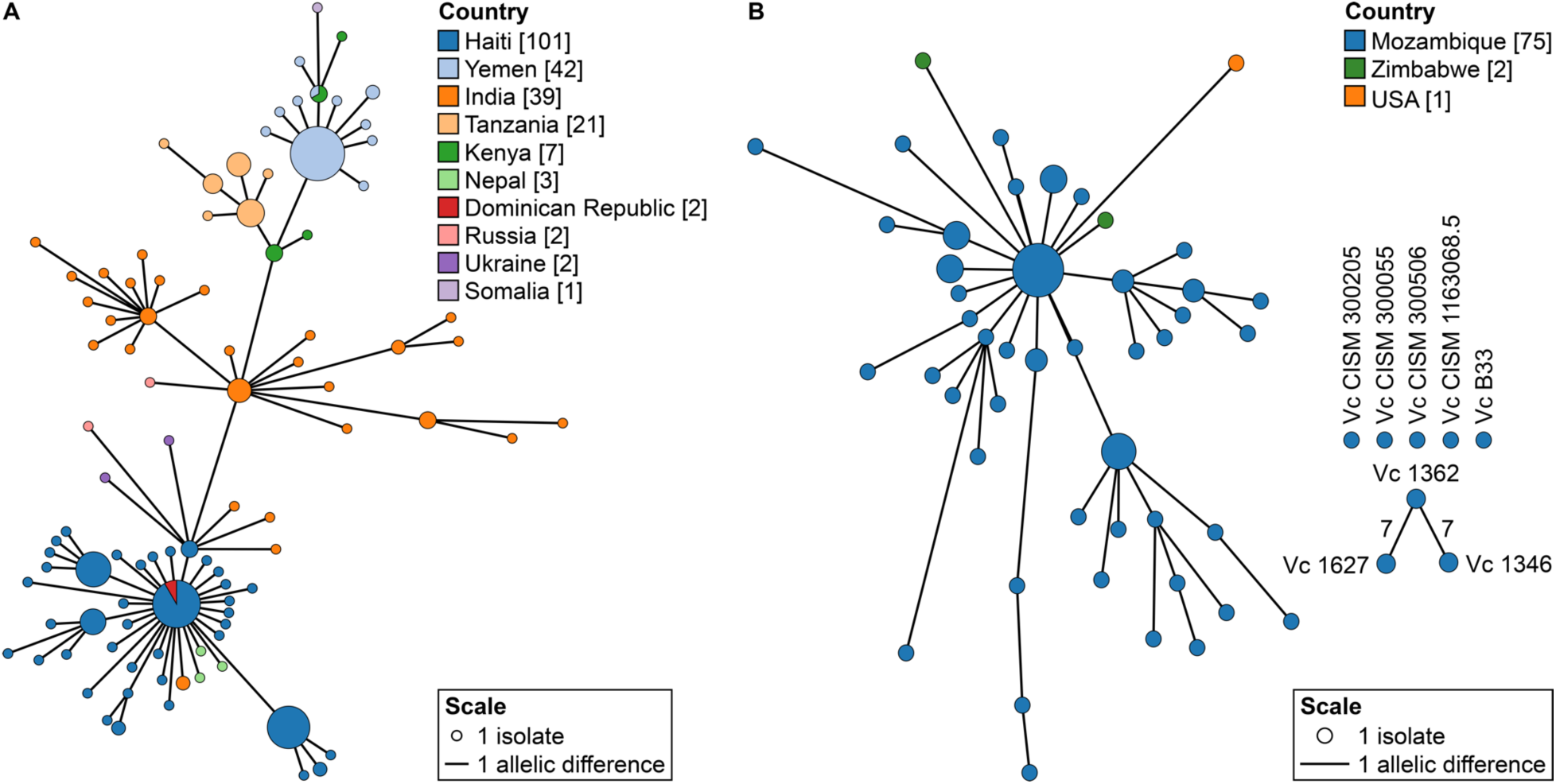
Minimum spanning trees isolated when the outbreak threshold was applied to the complete dataset of 1,262 isolates. A) All isolates that clustered together with the Haiti (dark blue) and Yemen (light blue) isolates based on the clustering threshold of seven allelic differences. B) All isolates that clustered with the Mozambique isolates (dark blue) based on the clustering threshold of seven allelic differences. Additional Mozambique isolates that are not part of the same outbreak cluster are also shown. Three isolates, two from Zimbabwe (green) and one from the USA (orange), are connected as they share seven or fewer allelic differences with the Mozambique isolates. In both panels, the size of the nodes is proportional to the number of isolates. Length of the lines is proportional to the number of allelic differences and all connections have less than or equal to seven allelic differences.

Cholera is still endemic in Africa (10) and caused several major reported outbreaks in different countries over last few decades (66) including Mozambique (23, 67) and Zimbabwe (68). Another major outbreak cluster groups most of the Mozambique isolates together with two Zimbabwe isolates (strains CP1038(11) and 2011EL-1137) and one USA isolate (2009V-1116) (Fig. 5B). Based on cgMLST, it is evident that the two Zimbabwe isolates are closely related to the Mozambique isolates, differing at only four or less alleles. The close proximity of the two countries suggests that these are likely travel-associated cases. Although outbreaks involving the Mozambique isolates (23) and the Zimbabwe isolates (58, 69) have been independently studied, the link between these isolates have not been shown before. Global cgMLST analysis is therefore an invaluable tool as it allows for the identification of links between independent studies. However, with only two Zimbabwe isolates in the dataset, additional sampling in this region is required to understand the epidemiology of this outbreak. According to the NCBI BioSample database, strain 2009V-1116 was collected by the Centers for Disease Control and Prevention in 2009 and is associated with travel to Pakistan. Since the 7^th^ pandemic El Tor lineage has been circulating in Asian and Middle Eastern countries for a long time (70), it is possible that, at least within our dataset, the Mozambique isolates are the closest relative to this specific Pakistan strain.

### Confirmation of an African connection for the Yemen outbreak

The Yemen cholera outbreak began in October 2016 with eleven confirmed cases (http://www.emro.who.int/pandemic-epidemic-diseases/cholera/cholera-cases-in-yemen.html). By January 2017, there were already over 10,000 cholera cases with 99 associated deaths (http://www.emro.who.int/pandemic-epidemic-diseases/cholera/weekly-update-cholera-cases-in-yemen-15-jan-2017.html). By the end of that year, there were over 900,000 cholera cases (http://www.emro.who.int/pandemic-epidemic-diseases/cholera/outbreak-update-cholera-in-yemen-19-december-2017.html). The outbreak continues on today as the largest cholera outbreak in modern history. As isolates from this outbreak were only recently made available (9), they were not part of the initial dataset for the cgMLST scheme development. These isolates were added and analyzed on PubMLST after the scheme had been established. This set of isolates therefore serves as an independent test of the universality and applicability of the cgMLST scheme. To determine the potential origin of the Yemen outbreak and its phylogenetic relationships with existing *V. cholerae* strains, the Yemen isolates were compared with other 7^th^ pandemic El Tor isolates from Asian and African countries (Table S3). All allele designations and cgST assignments were done automatically on PubMLST. MST was built using these isolates and all connections with seven and fewer allelic differences are represented as solid lines (Fig. 6). Isolates connected by solid lines therefore belong in the same outbreak cluster as defined by the outbreak threshold of seven allelic differences. Isolates from Yemen, Kenya, and Haiti all cluster with the central Indian isolates with seven or fewer allelic differences; however, the closest relatives to the Yemen isolates are those from Kenya with four or fewer allelic differences (Fig. 6). The Indian isolates are the next closest connection but there is no direct linkage between these and the Yemen isolate. This pattern is consistent with the work of Weill and colleagues (9), where they suggested that the Yemen outbreak strains may have come from East Africa which itself came from South Asia based on SNP-base phylogenetic analysis and Bayesian evolutionary analysis.

**Fig. 6.**
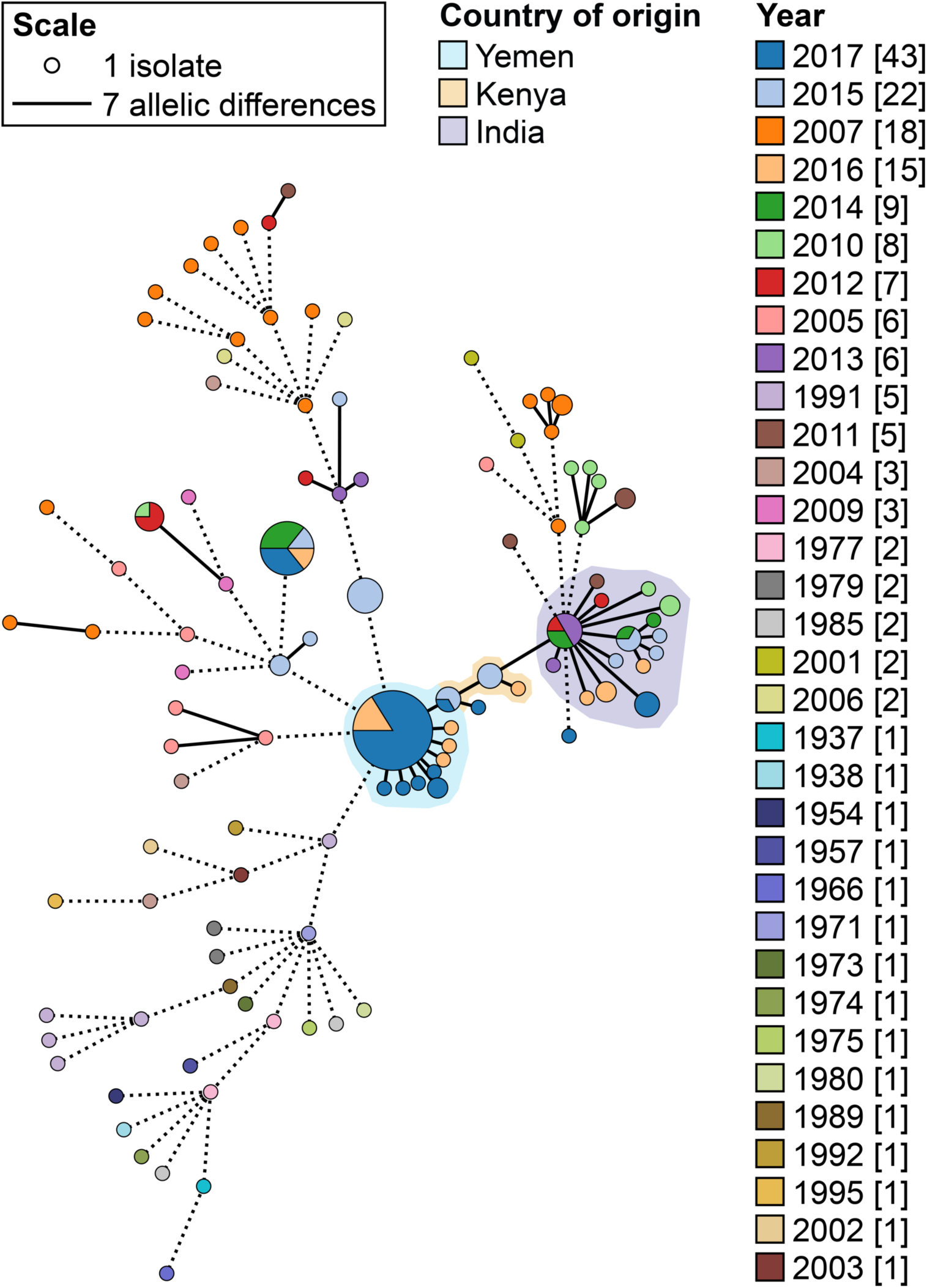
cgMLST MST of all Yemen isolates and representative 7^th^ pandemic El Tor strains. All isolates connected by dotted lines share eight or more allelic differences (not drawn to scale). All isolates connected with solid lines share seven or fewer allelic differences (i.e., they belong to the same outbreak cluster; drawn to scale). Each node represents a cgST that is colored by year of collection. The outbreak clusters are shaded by country.

Unlike the limited samples available from African cholera outbreaks, the Haiti and Yemen outbreaks are significant cases for epidemiological investigations because *V. cholerae* has been heavily sampled from these countries as well as surrounding regions. Two major limitations in genomic epidemiology have been the lack of a universal classification scheme and a comprehensive database; however, this is no longer the case in the genomic era as sequencing technology is becoming increasingly more accessible (8). A genomic approach, as shown here, is able to produce accurate predictions of potential origins of outbreaks and provides us with sufficient resolution to accurately track the spread of the disease. Therefore, genomic analysis should be the first step in any epidemiological study as not only will it help guide subsequent analyses and investigations, but consistently sequencing new genomes will also help expand and refine the current global *V. cholerae* genome database.

### Increased resolution for the history of cholera in Mozambique: comparing cgMLST to MLVA

The 7^th^ pandemic reached Africa in 1970 and cholera appeared in Mozambique at roughly the same time (57). Since its introduction, cholera has been endemic in that country and has continued to cause multiple outbreaks (23). A popular tool for outbreak investigation is MLVA (32, 38), which was recently used to study *V. cholerae* strains collected in Mozambique over multiple years (23). MLVA is a subspecies typing method similar to MLST in concept; however, it utilizes VNTRs instead of using gene sequences. As VNTR mutates at a faster rate than conserved genes, it has been shown that MLVA provides greater resolution than MLST for some species (32, 33). To establish a direct comparison between our cgMLST scheme and this MLVA scheme, we examined the MSTs created by both methods focusing only on shared isolates (Fig. 7A and 7B). The MLVA identified 26 profiles forming two clonal complexes (CCs) and four singletons (Fig. 7A) (23). A similar population structure is seen with the cgMLST analysis (Fig. 7B), including the four singletons identified in the MLVA. The central node in the cgMLST MST consists mostly of isolates with MLVA profile ‘8,4,6,18,21’ similar to the central node in the MLVA MST (23). The two CCs identified in the MLVA MST are also identified in the cgMLST MST with the smaller CC2 being at least four allelic differences away from the larger CC1.

**Fig. 7.**
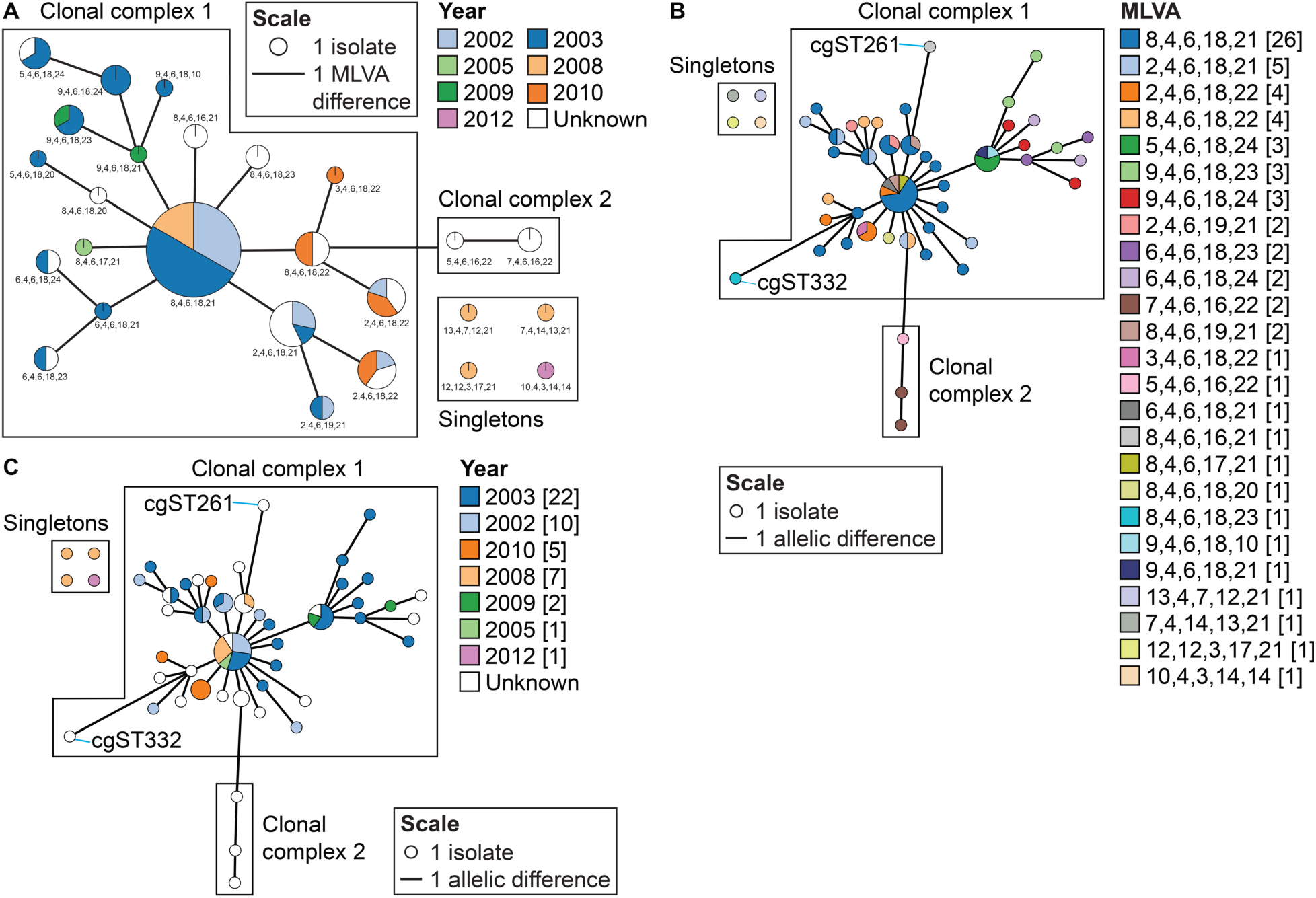
Comparison between cgMLST and MLVA with a focus on the Mozambique isolates. A) Population structure of pandemic *V. cholerae* in Mozambique based on MLVA profiles by Garrine and colleagues (23). MST of the Mozambique isolates based on the cgMLST scheme colored based on B) MLVA profiles and C) year of isolation. All isolates in B and C connected with lines share seven or fewer allelic differences. For all panels, the size of the nodes is proportional to the number of isolates. The length of the lines is proportional to the number of allelic differences.

Although there are a few MLVA types that were grouped into a single cgST, such as cgST1 and cgST114, indicating that cgMLST was unable to resolve the differences in these MLVA types, there are many MLVA types such as profile ‘2,4,6,18,21’, profile ‘7,4,6,16,22’, profile ‘9,4,6,18,24’, and profile ‘8,4,6,18,22’ that were split into multiple cgSTs. Overall there are 48 cgSTs as opposed to only 26 MLVA types, showing that cgMLST provides better resolution overall than MLVA. The cgMLST analysis overlaid with isolation dates shows that the Mozambique *V. cholerae* strains are highly clonal and strains from the same cgST can cause outbreaks over multiple years (e.g., cgST114 and cgST94) (Fig. 7C), which corroborates the claim made in the initial MLVA study that the same MLVA type can be seen over multiple years (23). In addition to increased resolution, cgMLST also produces more reliable and reproducible results than MLVA, as it eliminates errors associated with the detection of VNTR regions using PCR or sequencing-based methods. For the same reason that MLST is less affected by convergent evolution compared to MLVA (35), cgMLST is also less affected by convergent evolution.

### Standardizing the genotypes responsible for the Haiti 2010 cholera outbreak: comparing cgMLST and SNP-based analyses

One of the largest cholera outbreaks in modern history occurred in Haiti following the devastating earthquake in 2010 (8, 71). Prior to this outbreak, there were no documented cholera cases in Haiti (14, 18). Since the initial introduction, *V. cholerae* now remains endemic in Haiti and is responsible for thousands of cholera cases annually (71). Multiple studies have strongly suggested that the Haitian strains were in fact imported from Nepal (by the UN Nepalese troops) and the outbreak occurred as a result of both inappropriate sanitary practice and the lack of screening of the UN troops upon their arrival in Haiti (8, 15, 16, 71).

A SNP-based approach was used to study the evolutionary dynamics of *V. cholerae* in Haiti (14). This technique relies on the identification of SNPs in draft or closed genomes. The primary benefit of this method is that assembly and annotation are not required. It is also capable of resolving closely related strains using whole genome data. However, SNP-based methods are highly influenced by recombination events (43, 44) and quality filter parameters chosen (72).

To establish a direct comparison between the cgMLST scheme and SNP-based analysis, we focused on MSTs of only the Haitian outbreak isolates (Fig. 8). All Haitian isolates are closely related according to the cgMLST scheme, sharing at most four allelic differences with each other (Fig. 8A). The Haitian and Nepalese isolates, therefore, also belong to the same sublineage (SL6) which is consistent with the fact that these isolates belong to the same MLST ST (either ST1 or ST69 based on the 2016 or 2013 MLST scheme, respectively (20, 21)) (Table S1). The overall population structure is similar between the two methods where we have SNP ST1 as the center of the MST with ST2 and ST3 extending from that likely ancestral genotype (Fig. 8). SNP ST1, ST2, and ST3 can be split into 11, 2, and 3 different cgSTs, respectively (Fig. 8A). There is only one case, that of cgST66, where it contains isolates from both SNP ST1 and ST3. Overall, cgMLST was able to differentiate 39% of the isolates while the SNP-based analysis differentiated 35%, showing comparable level of resolution. As expected, both the cgMLST and the SNP-based analyses showed that the Haiti outbreak is highly clonal where most isolates belong to the same cgST or SNP ST (14). However, one important advantage of cgMLST over SNP-based analysis is that the former can be easily standardized because it relies on a predefined set of core genes. Based on these standardized genes, a systematic nomenclature system can be established. This makes cgMLST more suitable than the SNP-based method as a universally applicable classification system for epidemiological studies and research worldwide.

**Fig. 8.**
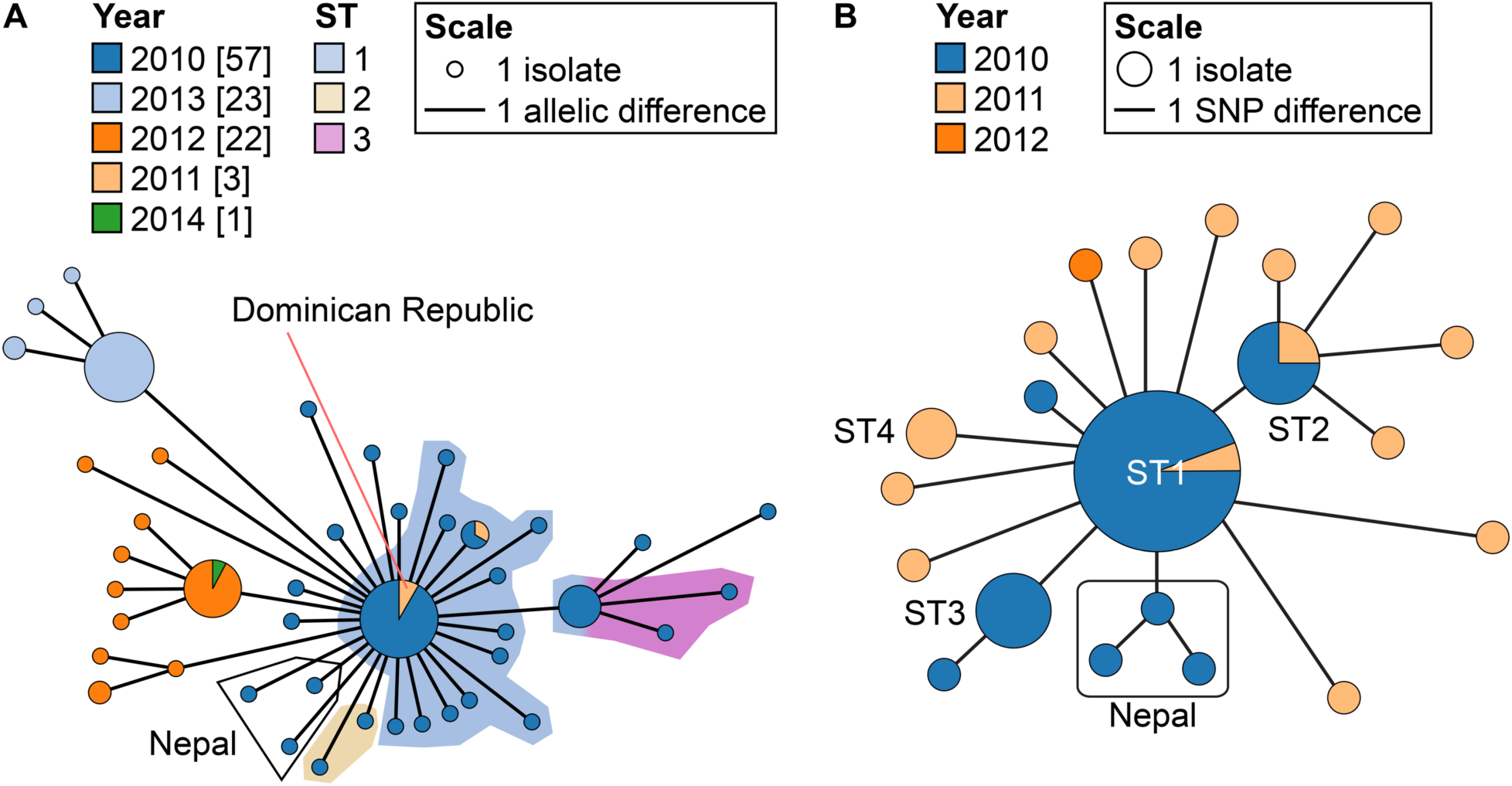
Comparison between cgMLST and SNP-based analysis with a focus on the Haiti outbreak and related isolates. A) MST of isolates from the 2010 cholera outbreak in Haiti. All lines indicate connections of four or fewer allelic differences. Each node represents a cgST which is colored by year of isolation. Background shading represent ST designations based on 45 high-quality SNPs by Katz and colleagues (14). Note that cgST66 contains a mix of color as it contains both ST1 and ST3. Any isolate from countries other than Haiti is indicated. The length of the lines is proportional to the number of allelic differences. B) MST constructed from whole genome SNP data (14). The length of the lines indicates the number of nucleotide substitutions. Size of the nodes is proportional to the number of isolates.

### Environmental isolates differ from clinical strains by their diversity and their associations with specific geographical locations

To look at the geographic signal of *V. cholerae*, we eliminated all clinical isolates and those that belong to the PG lineage (18, 19). This is because the geographic signal of clinical strains can be skewed, as pathogenic strains can travel long distances in a short period of time through association with human hosts. The geographical analysis was therefore performed only with environmental isolates.

Along with all the publicly available environmental strains that are not part of the PG lineage, there are a total of 195 isolates spanning 9 countries. After grouping the isolates at the sublineage level (i.e., each cluster have at most 133 allelic differences), it could be noted that all isolates from the same sublineage also shared a country of origin, with the exception of strains 692-79 and 857 (Fig. 9), which are from the USA and Bangladesh, respectively. Phylogenetic analysis shows these isolates to be closely related to strain A215, a clinical isolate from the USA (Fig. 4). All three strains contain the *toxR* gene, a toxin transcriptional regulator common in pathogenic *V. cholerae* (73), as well as genes encoding for the Mannose-sensitive hemagglutinin pilus, the RTX toxin, and hemolysin (*hlyA*), all of which are putative virulence factors for this species. In addition, strains A215 and 857 also harbor the zona occludens toxin gene. Similar toxin gene contents among these three isolates and close phylogenetic relationships suggest that strains 692-79 and 857 may also be potentially pathogenic and capable of surviving inside a human host. This provides evidence that although clinical isolates can spread across the world rapidly and closely related isolates can be from very different parts of the world, environmental isolates from the same geographic origin share an affinity among each other at least at the sublineage level. It is important to note that our current dataset contains a relatively small number of environmental isolates that are not part of the PG lineage. Therefore, this distinct distribution pattern based on geographic origin may be a result of currently insufficient sampling of environmental *V. cholerae* worldwide. With large-scale environmental sampling, it will be possible to determine with greater accuracy the evolutionary rate and distribution pattern of *V. cholerae* in the environment using cgMLST. In addition, this method will become an invaluable tool in dealing with these large datasets, as it provides an efficient and standardized method of classification.

**Fig. 9.**
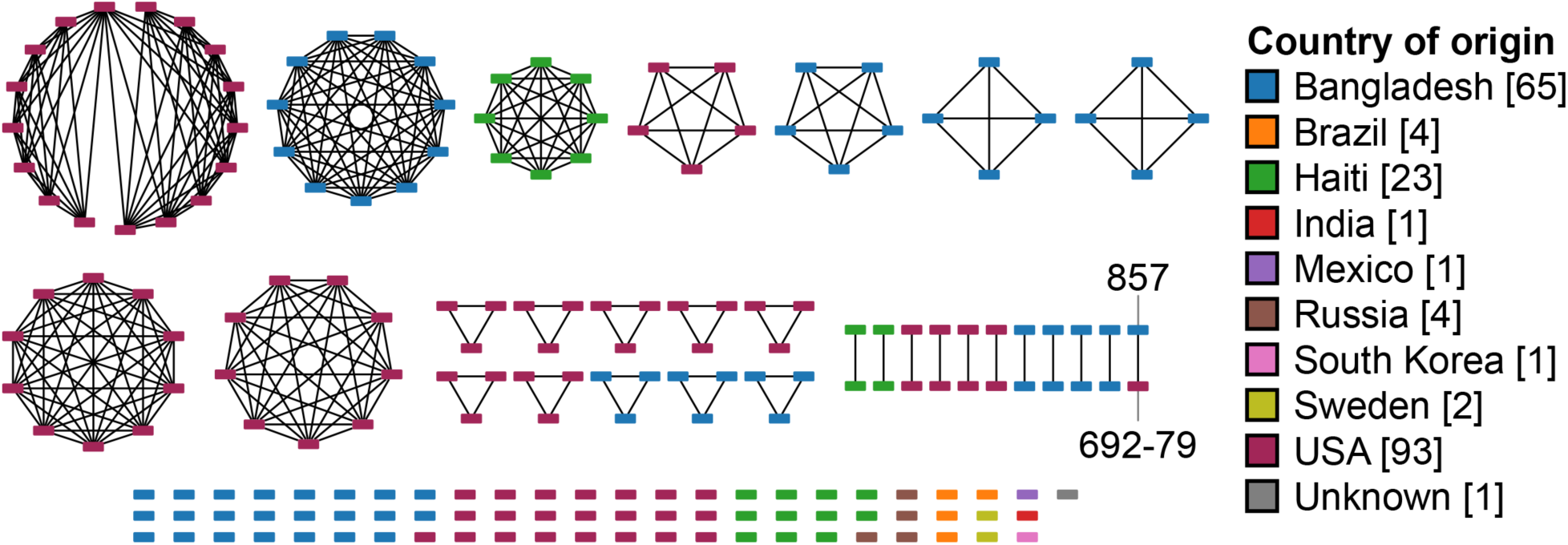
Sublineage clusters of non-clinical environmental isolates that are not part of the PG lineage. Clusters are constructed using networkX (95) and visualized with Cytoscape (96). Missing loci were assumed to contain the most common allele when calculating allelic differences. Isolates are connected only if they share 133 allelic differences or fewer with each other. Each node represents an isolate and is colored by the country of origin.

### Conclusion

With an extensive collection of over 1,200 *V. cholerae* isolates, we developed a cgMLST scheme based on 2,443 core genes. We established a sublineage-level definition based on 133 allelic differences as part of our standardized classification scheme. It was determined by comparisons with previous MLST schemes that the cgMLST sublineage classification can be used as a proxy for traditional MLST. Additionally, the universality and applicability of the scheme have been tested by looking at various cholera outbreak cases. We determined an outbreak threshold based on seven allelic differences that groups isolates from the same outbreak together with strains from the potential source of introduction. This threshold creates clusters that are consistent with known epidemiological data when applied to the Haiti and Yemen cholera outbreaks, two of the best-documented cholera outbreaks in modern history. Also, we were able to confirm the South Asian origin of modern cholera outbreaks. Furthermore, although current sampling is limited, a geographic signal at the sublineage level not seen in clinical strains could be identified among environmental isolates that are not part of the PG lineage (18, 19). Lastly, this scheme is fully implemented on PubMLST (https://pubmlst.org/vcholerae/) for public access. All newly available genomes uploaded to PubMLST will be annotated automatically and a cgST designation will be assigned to isolates with less than 100 missing loci. Relevant epidemiological data and the variety of analytical and visualization tools are all integrated on PubMLST, allowing for a quick analysis of any newly sequenced genome in a global context. This scheme will be an important tool for future large-scale epidemiological and biogeographical research.

## MATERIALS AND METHODS

### Dataset description

On November 6^th^ 2018, 1,172 *V. cholerae* genomes consisting of 800 draft and complete genomes and 372 sequence read archives (SRAs), available from both publicly available database and private collection, were selected as our dataset. One hundred sixteen SRAs from a recent study on the Yemen cholera outbreak (9) were subsequently added as an independent evaluation of the cgMLST scheme (Table S3). The 488 SRAs were assembled using skesa (74) or the CLC Genomics Workbench 7 (QIAGEN) using default parameters. This total dataset of 1,288 included twenty-six genomes with less than 90% of the core genes, which were identified using USearch (75) based on RAST (76) annotations. These 26 genomes were removed from subsequent analyses resulting in a final dataset of 1,262 genomes collected from 52 countries and spanning 82 years from 1937 to 2018 (Table S1). These include a historical collection from the 6^th^ cholera pandemic, clinical isolates from outbreaks in various countries (e.g., Bangladesh, India, Haiti, Yemen, the Democratic Republic of Congo, and Russia), and environmental isolates from different parts of the world (e.g., Bangladesh, Haiti, USA, Mexico, Brazil, etc.).

### Gene identification and allele assignments

Instead of using the full dataset of 1,288 genomes, we selected a subset of high-quality genomes because core gene identification is highly dependent on the initial dataset and the inclusion of poorly assembled and/or sequenced data will reduce the number of core genes identified (49). Firstly, 800 already assembled draft or complete genomes were selected for core gene identification. Low-quality assemblies were then eliminated by removing genomes with less than 40× coverage and/or N50 values less than 40 kb. From a previously established cgMLST scheme for *L. monocytogenes*, 40× coverage and 20 kb N50 value were used as cutoff thresholds, as genomes that do not meet these criteria resulted in a low proportion of loci being called (49). The 40× coverage cutoff was adopted for this study; however, because the average *V. cholerae* genome size (∼ 4 Mb) is larger than the average *L. monocytogenes* genome (∼ 3 Mb), 40 kb was instead selected as the N50 cutoff. The use of these cutoffs resulted in the removal of 82 genomes.

The remaining 718 genomes were annotated using RAST (76) and USearch (75), and a tentative set of core genes that were on average present in 99% of the genomes were selected. An additional 13 genomes were removed, as they lacked more than 90% for the core genes, leaving us with a dataset of 705 high-quality genomes (Table S4). However, an additional 26 genomes were subsequently removed for the core gene analysis as it has been previously suggested that they form a highly divergent lineage within the *V. cholerae* (77–79), ensuring that the dataset used for core gene identification consists only of unambiguously *V. cholerae* isolates (also as verified by average nucleotide identity (80) and digital DNA–DNA hybridization (81) between genomes (77–79)). Completeness and potential contamination of all remaining 679 genomes were also independently evaluated by checkM, which estimates these values based on the presence and number of copies of a set of pre-defined single copy marker genes (82) (Table S5). All genomes were, according to the criteria established by checkM, nearly complete (≥97%) with medium to low levels of contamination (<7%) (82).

Each orthologous gene was compared against the *V. cholerae* N16961 reference genome using BLASTN (83) to determine gene function. Any gene family with no homolog in N16961 or classified as pseudogenes on the NCBI GenBank database were removed, meaning N16961 was 100% complete for the cgMLST scheme. Any gene that was present in more than one copy in any of the initial 679 genomes was also removed, as they were considered paralogous. Thus, in this context, core genes are defined as being present in at least 90% of the 679 high-quality assembled genomes in a single copy. By choosing a relaxed cutoff of 90% completeness, we accounted for missing genes due to sequencing, annotation, or assembly errors while ensuring there is sufficient resolution to differentiate between closely related strains, with at least 2,199 loci remaining for classification purposes. The final cgMLST scheme utilizes a set of 2,443 single-copy core gene loci, which is 2,425,296 bp in size and covering approximately 61% of the genome. The list of core genes is available on PubMLST (https://pubmlst.org/vcholerae/).

Automated scripts in BIGSdb (84) were used to perform allele calls and assignments for all 1,262 isolates (Table S1). Allele calls were made only for complete coding sequences with a minimum of 70% similarity and 70% length coverage at the nucleotide level, as previously described (49). Default settings were used for all other parameters.

### Core genome sequence type (cgST) assignment

cgST, which was defined as a unique combination of alleles of all loci included in the scheme, was assigned for all isolates, excluding those from the Yemen outbreak study (8), with an in-house script, as previously described (85). Briefly, missing loci were replaced with the most common allele when assigning cgSTs, allowing for a conservative estimate of diversity (85). The 116 isolates from the Yemen cholera outbreak study (9) were annotated automatically by uploading them to PubMLST. PubMLST treated missing alleles as ‘N’. cgSTs were assigned to each allele profile, treating ‘N’ as a regular allele designation. However, different from typical allele designations, ‘N’s can represent any allelic sequence; therefore, some isolates may contain multiple cgST designations, all of which are possibly true cgSTs. For isolates with more than one cgST suggested by PubMLST, postprocessing was done using an in-house script to identify the most likely cgST, which was determined by assuming missing loci contained the most common allele (Table S2). It is expected that as genome sequencing becomes more reliable, higher quality genomes will be available and any missing data can be updated as needed.

### MLST scheme and sequence type (ST) assignments

Two MLST schemes developed for *V. cholerae* were mapped to this cgMLST scheme. The first MLST scheme developed in 2013 by Octavia and colleagues (20) was used to study the global population structure of non-O1/non-O139 *V. cholerae* and is currently hosted on PubMLST. All isolates uploaded to PubMLST were automatically annotated with this scheme. Any missing data in this scheme was ignored and no ST designation was assigned. The second MLST scheme developed in 2016 by Kirchberger and colleagues (21) was used to study the population structure of environmental *V. cholerae* in a region on the US East Coast. The second MLST scheme is not currently hosted on PubMLST, but because the housekeeping genes in this scheme are also found in the cgMLST scheme, a similar in-house script used in cgST assignments was used to assign ST designations. Therefore, all isolates in this study were assigned three designations when possible – two ST designations based on the two previously established MLST schemes (20, 21) and one cgST designation based on the cgMLST scheme from this study.

### Outbreak and sublineage clustering thresholds

A clustering threshold was defined as the maximum number of allelic differences found within a cluster. All clusters were produced based on the single-linkage clustering method, which meant an isolate belonged to a cluster if it linked with any isolate within that cluster. Two metrics were used as general guidelines for determining clustering thresholds. The first metric used was the Dunn Index (DI), which measured clustering efficiencies (60). Briefly, the DI was highest for a network (i.e., the network has the best clustering efficiency) when the intra-cluster distances were minimized, and the inter-cluster distances were maximized. Since isolate distances were measured based on allelic differences, a high DI resulted in clusters where isolates were more closely related to those found within the same cluster than those found in a different cluster. The DI was calculated using the R packages ‘clvalid’ and ‘boot’ with 100 bootstrap replicates for each threshold and graphed using the R package ‘ggplot2’ (86–89).

The second metric used was the Adjusted Rand Index (ARI), which measured the level of similarity between two networks when clustering the same set of isolates by measuring the amount of agreements (i.e., the number of pairs that were grouped either as being in the same cluster or different cluster in both networks) and disagreements (i.e., the number of pairs that were grouped together in one network but grouped separately in another) (62). The values ranged from −1 (i.e., two networks are exactly opposite) to 1 (i.e., two networks are identical). ARI was used to determine the level of similarity between various clustering thresholds and the MLST schemes. ARI was calculated using the R package ‘clues’ and graphed using ‘ggplot2’ (87, 89, 90).

### Minimum spanning tree (MST)

All MSTs, unless otherwise specified, were constructed using GrapeTree MSTv2 (91). Loci with missing data were included in the profile as “–”. GrapeTree provided a novel algorithm that accounted for missing data when constructing an MST, an important feature since missing data is common in whole and core genome-based analyses. GrapeTree is currently integrated within PubMLST, which allows for quick visualization of the dataset with any provenance data.

### Phylogenetic analysis

Parsnp v1.2 (92) was used to reconstruct the phylogenetic tree using *V. cholerae* N16961 as the reference genome. The -x flag was used to enable filtering of SNPs in recombinogenic regions as identified by PhiPack (93). Default settings were used for all other parameters. The phylogenetic tree included 1,146 genomes (all genomes except for the 116 isolates from the recent Yemen cholera outbreak study (9)). Since all isolates sequenced for the latter study belonged to the 7^th^ pandemic El Tor lineage, it would have had limited impact on the overall structure of the tree. The phylogeny was visualized and annotated using iTOL (94).

### Biogeographical analysis of environmental isolates

All environmental isolates that were not part of the PG lineage (18, 19) were first clustered based on the sublineage threshold using the python package ‘networkX’ (95). Missing alleles were replaced with the most common allelic designation when calculating pairwise differences to establish a more conservative estimate of diversity. The network was then visualized using Cytoscape (96).

### Data availability

All previously sequenced *V. cholerae* genomes and the additional 65 genomes sequenced in this study are available on the NCBI GenBank database. Table S6 lists all the accession numbers for all the genomes used in this study. In addition, all genome sequences, allelic profiles, cgST designations, ST designations, and relevant epidemiological data are publicly available on PubMLST (https://pubmlst.org/vcholerae/).

## ACKNOWLEDGMENTS

KYHL, FDO, and YFB conceived the experiments. KYHL, FDO, and MTI performed all data collection and analyses. FDO and TN performed genome sequencing of Bangladesh isolates. MA and CLT provided isolates used in this study.

We thank Dr. Keith Jolley (University of Oxford) for providing valuable feedback regarding the development of the cgMLST scheme, as well as the implementation of this scheme on PubMLST. We also thank Monica Im (Centers for Disease Control and Prevention) for assistance with obtaining whole genome sequences.

This work was supported by the Natural Sciences and Engineering Research Council (NSERC) of Canada (to YFB); the Integrated Microbial Biodiversity program of the Canadian Institute for Advanced Research (to YFB); federal appropriations to the Centers for Disease Control and Prevention through the Advanced Molecular Detection Initiative (to CLT); and graduate student scholarships from Alberta Innovates – Technology Futures (to KYHL, FDO, MTI, and TN), NSERC (to KYHL and TN), the University of Alberta Faculty of Graduate Studies and Research (Queen Elizabeth II Graduate Scholarship to KYHL and TN), and the Bank of Montréal Financial Group (to FDO). MA of icddr,b acknowledges the governments of Bangladesh, Canada, Sweden, and the United Kingdom for providing core/unrestricted support. The funders had no role in study design, data collection and interpretation, or the decision to submit the work for publication. The findings and conclusions in this report are those of the authors and do not necessarily represent the official position of the Centers for Disease Control and Prevention.

## REFERENCES

1. Jahan S. 2016. Cholera - epidemiology, prevention and control, p. 145–157. In Makun, HA (ed.), Significance, Prevention and Control of Food Related Diseases. Croatia: InTech. InTechOpen, Rijeka, Croatia.

2. Momba M, Azab El-Liethy M. 2017. *Vibrio cholerae* and Cholera biotypes, p. online. *In* Pruden, A, Ashbolt, N, Miller, J (eds.), Global Water Pathogen Project. Michigan State University, Michigan.

3. Clemens JD, Nair GB, Ahmed T, Qadri F, Holmgren J. 2017. Cholera. Lancet 390:1539–1549.

4. Kaper JB, Morris JG, Levine MM. 1995. Cholera. Clin Microbiol Rev 8:48–86.

5. Islam MT, Alam M, Boucher Y. 2017. Emergence, ecology and dispersal of the pandemic generating Vibrio cholerae lineage. Int Microbiol 20:106–115.

6. Choi SY, Rashed SM, Hasan NA, Alam M, Islam T, Sadique A, Johura F-T, Eppinger M, Ravel J, Huq A, Cravioto A, Colwell RR. 2016. Phylogenetic Diversity of *Vibrio cholerae* Associated with Endemic Cholera in Mexico from 1991 to 2008. MBio 7:e02160–15.

7. Dalsgaard A, Skov MN, Serichantalergs O, Echeverria P, Meza R, Taylor DN. 1997. Molecular evolution of *Vibrio cholerae* O1 strains isolated in Lima, Peru, from 1991 to 1995. J Clin Microbiol 35:1151–1156.

8. Orata FD, Keim PS, Boucher Y. 2014. The 2010 Cholera Outbreak in Haiti: How Science Solved a Controversy. PLoS Pathog 10:e1003967.

9. Weill F-X, Domman D, Njamkepo E, Almesbahi AA, Naji M, Nasher SS, Rakesh A, Assiri AM, Sharma NC, Kariuki S, Pourshafie MR, Rauzier J, Abubakar A, Carter JY, Wamala JF, Seguin C, Bouchier C, Malliavin T, Bakhshi B, Abulmaali HH., Kumar D, Njoroge SM, Malik MR, Kiiru J, Luquero FJ, Azman AS, Ramamurthy T, Thomson NR, Quilici M-L. 2018. Genomic insights into the 2016–2017 cholera epidemic in Yemen. Nature 565:230–233.

10. Ali M, Nelson AR, Lopez AL, Sack DA. 2015. Updated global burden of cholera in endemic countries. PLoS Negl Trop Dis 9:e0003832.

11. Legros D. 2018. Global cholera epidemiology: Opportunities to reduce the burden of cholera by 2030. J Infect Dis 218:S137–S140.

12. Mintz E, Omolo J, Ope M, Gathigi L, Thuranira M, Loharikar A, Abade A, Ayers T, De Cock KM, Makayotto L, Oundo J, Ismail AM, Breiman RF, Langat D, Briere E, Amwayi S, O’Reilly CE, Njeru I. 2013. A National Cholera Epidemic With High Case Fatality Rates-Kenya 2009. J Infect Dis 208:S69–S77.

13. Sack DA, Sack RB, Chaignat C-L. 2006. Getting Serious about Cholera. N Engl J Med 355:649–651.

14. Katz LSS, Petkau A, Beaulaurier J, Tyler S, Antonova ESS, Turnsek MAA, Guo Y, Wang S, Paxinos EEE, Orata F, Gladney LMM, Stroika S, Folster JPP, Rowe L, Freeman MMM, Knox N, Frace M, Boncy J, Graham M, Hammer BKK, Boucher Y, Bashir A, Hanage WPP, Domselaar GV Van, Tarr L, Van Domselaar G, Tarr CLL, Domselaar GV Van. 2013. Evolutionary Dynamics of *Vibrio cholerae* O1 following a Single-Source Introduction to Haiti. MBio 4:e00398–13.

15. Frerichs RR. 2016. Deadly river: Cholera and cover-up in post-earthquake Haiti. Cornell University Press, Ithaca.

16. Frerichs RR, Keim PS, Barrais R, Piarroux R. 2012. Nepalese origin of cholera epidemic in Haiti. Clin Microbiol Infect 18:E158–E163.

17. Hendriksen RS, Price LB, Schupp JM, Gillece JD, Kaas RS, Engelthaler DM, Bortolaia V, Pearson T, Waters AE, Prasad Upadhyay B, Devi Shrestha S, Adhikari S, Shakya G, Keim PS, Aarestrup FM. 2011. Population Genetics of *Vibrio cholerae* from Nepal in 2010: Evidence on the Origin of the Haitian Outbreak. MBio 2:e00157–11.

18. Boucher Y. 2016. Sustained Local Diversity of *Vibrio cholerae* O1 Biotypes in a Previously Cholera-Free Country. MBio 7:e00570–16.

19. Chun J, Grim CJ, Hasan NA, Lee JH, Choi SY, Haley BJ, Taviani E, Jeon Y-S, Kim DW, Lee J-H, Brettin TS, Bruce DC, Challacombe JF, Detter JC, Han CS, Munk AC, Chertkov O, Meincke L, Saunders E, Walters RA, Huq A, Nair GB, Colwell RR. 2009. Comparative genomics reveals mechanism for short-term and long-term clonal transitions in pandemic *Vibrio cholerae*. Proc Natl Acad Sci 106:15442–15447.

20. Octavia S, Salim A, Kurniawan J, Lam C, Leung Q, Ahsan S, Reeves PR, Nair GB, Lan R. 2013. Population Structure and Evolution of Non-O1/Non-O139 *Vibrio cholerae* by Multilocus Sequence Typing. PLoS One 8:e65342.

21. Kirchberger PC, Orata FD, Barlow EJ, Kauffman KM, Case RJ, Polz MF, Boucher Y. 2016. A small number of phylogenetically distinct clonal complexes dominate a coastal *Vibrio cholerae* population. Appl Environ Microbiol 82:5576–5586.

22. Bwire G, Sack DA, Almeida M, Li S, Voeglein JB, Debes AK, Kagirita A, Buyinza AW, Orach CG, Stine OC. 2018. Molecular characterization of *Vibrio cholerae* responsible for cholera epidemics in Uganda by PCR, MLVA and WGS. PLoS Negl Trop Dis 12:e0006492.

23. Garrine M, Mandomando I, Vubil D, Nhampossa T, Acacio S, Li S, Paulson JN, Almeida M, Domman D, Thomson NR, Alonso P, Stine OC. 2017. Minimal genetic change in *Vibrio cholerae* in Mozambique over time: Multilocus variable number tandem repeat analysis and whole genome sequencing. PLoS Negl Trop Dis 11:e0005671.

24. Safa A, Nair GB, Kong RYC. 2010. Evolution of new variants of *Vibrio cholerae* O1. Trends Microbiol 18:46–54.

25. Boucher Y, Orata FD, Alam M. 2015. The out-of-the-delta hypothesis: Dense human populations in low-lying river deltas served as agents for the evolution of a deadly pathogen. Front Microbiol 6:1120.

26. Mandal S, Mandal MD, Pal NK. 2011. Cholera: A great global concern. Asian Pac J Trop Med 4:573–580.

27. Maiden MCJ, Bygraves JA, Feil E, Morelli G, Russell JE, Urwin R, Zhang Q, Zhou J, Zurth K, Caugant DA, Feavers IM, Achtman M, Spratt BG. 1998. Multilocus sequence typing: A portable approach to the identification of clones within populations of pathogenic microorganisms. Proc Natl Acad Sci USA 95:3140–3145.

28. Horwood P, Collins D, Jonduo M, Rosewell A, Dutta S, Dagina R, Ropa B, Siba P, Greenhill A. 2011. Clonal Origins of *Vibrio cholerae* O1 El Tor Strains, Papua New Guinea, 2009–2011. Emerg Infect Dis 17:2063.

29. Luo Y, Ye J, Jin D, Ding G, Zhang Z, Mei L, Octavia S, Lan R. 2013. Molecular analysis of non-O1/non-O139 *Vibrio cholerae* isolated from hospitalised patients in China. BMC Microbiol 13:52.

30. Maiden MCJ, Van Rensburg MJJ, Bray JE, Earle SG, Ford SA, Jolley KA, McCarthy ND. 2013. MLST revisited: The gene-by-gene approach to bacterial genomics. Nat Rev Microbiol 11:728–736.

31. Gonzalez-Escalona N, Martinez-Urtaza J, Romero J, Espejo T R, Jaykus L-A, DePaola A. 2008. Determination of Molecular Phylogenetics of *Vibrio parahaemolyticus* Strains by Multilocus Sequence Typing. J Bacteriol 190:2831–2840.

32. Lam C, Octavia S, Reeves PR, Lan R. 2012. Multi-locus variable number tandem repeat analysis of 7th pandemic *Vibrio cholerae*. BMC Microbiol 12:82.

33. Chenal-Francisque V, Passet V, Brisse S, Cantinelli T, Diancourt L, Pourcel C, Lecuit M, Leclercq A, Tran-Hykes C, Bracq-Dieye H. 2013. Optimized Multilocus Variable-Number Tandem-Repeat Analysis Assay and Its Complementarity with Pulsed-Field Gel Electrophoresis and Multilocus Sequence Typing for *Listeria monocytogenes* Clone Identification and Surveillance. J Clin Microbiol 51:1868–1880.

34. Vogler AJ, Birdsell DN, Lee J, Vaissaire J, Doujet CL, Lapalus M, Wagner DM, Keim P. 2011. Phylogeography of *Francisella tularensis* ssp. holarctica in France. Lett Appl Microbiol 52:177–180.

35. Struelens MJ, Brisse S. 2013. From molecular to genomic epidemiology: Transforming surveillance and control of infectious diseases. Eurosurveillance 18:20386.

36. Sabat AJ, Budimir A, Nashev D, Sá-Leão R, van Dijl JM, Laurent F, Grundmann H, Friedrich AW, on behalf of the ESCMID Study Group. 2013. Overview of molecular typing methods for outbreak detection and epidemiological surveillance. Eurosurveillance 18:20380.

37. Klassen JL, Currie CR. 2012. Gene fragmentation in bacterial draft genomes: extent, consequences and mitigation. BMC Genomics 13:14.

38. Danin-Poleg Y, Cohen LA, Gancz H, Broza YY, Goldshmidt H, Malul E, Valinsky L, Lerner L, Broza M, Kashi Y. 2007. *Vibrio cholerae* Strain Typing and Phylogeny Study Based on Simple Sequence Repeats. J Clin Microbiol 45:736–746.

39. Wong VK, Baker S, Connor TR, Pickard D, Page AJ, Dave J, Murphy N, Holliman R, Sefton A, Millar M, Dyson ZA, Dougan G, Holt KE. 2016. An extended genotyping framework for *Salmonella enterica* serovar Typhi, the cause of human typhoid. Nat Commun 7:12827.

40. Leekitcharoenphon P, Nielsen EM, Kaas RS, Lund O, Aarestrup FM. 2014. Evaluation of Whole Genome Sequencing for Outbreak Detection of *Salmonella enterica*. PLoS One 9:e87991.

41. Chen C, Zhang W, Zheng H, Lan R, Wang H, Du P, Bai X, Ji S, Meng Q, Jin D, Liu K, Jing H, Ye C, Gao GF, Wang L, Gottschalk M, Xu J. 2013. Minimum core genome sequence typing of bacterial pathogens: A unified approach for clinical and public health microbiology. J Clin Microbiol 51:2582–2591.

42. Qin T, Zhang W, Liu W, Zhou H, Ren H, Shao Z, Lan R, Xu J. 2016. Population structure and minimum core genome typing of Legionella pneumophila. Sci Rep 6:21356.

43. Wang R, Yu D, Yue J, Kan B. 2016. Variations in SXT elements in epidemic *Vibrio cholerae* O1 El Tor strains in China. Sci Rep 6:22733.

44. Meibom KL, Blokesch M, Dolganov NA, Wu C-Y, Schoolnik GK. 2005. Chitin induces natural competence in *Vibrio cholerae*. Science 310:1824–1827.

45. Borgeaud S, Metzger LC, Scrignari T, Blokesch M. 2015. The type VI secretion system of *Vibrio cholerae* fosters horizontal gene transfer. Science 347:63–68.

46. Orata FD, Kirchberger PC, Méheust R, Barlow EJ, Tarr CL, Boucher Y. 2015. The Dynamics of Genetic Interactions between *Vibrio metoecus* and *Vibrio cholerae*, Two Close Relatives Co-Occurring in the Environment. Genome Biol Evol 7:2941–2954.

47. Boucher Y, Cordero OX, Takemura A, Hunt DE, Schliep K, Bapteste E, Lopez P, Tarr CL, Polz MF. 2011. Local Mobile Gene Pools Rapidly Cross Species Boundaries To Create Endemicity within Global *Vibrio cholerae* Populations. MBio 2.

48. Neumann B, Prior K, Bender JK, Harmsen D, Klare I, Fuchs S, Bethe A, Zühlke D, Göhler A, Schwarz S, Schaffer K, Riedel K, Wieler LH, Werner G. 2019. A Core Genome Multilocus Sequence Typing Scheme for *Enterococcus faecalis*. J Clin Microbiol 57:e01686–18.

49. Moura A, Criscuolo A, Pouseele H, Maury MM, Leclercq A, Tarr C, Björkman JT, Dallman T, Reimer A, Enouf V, Larsonneur E, Carleton H, Bracq-Dieye H, Katz LS, Jones L, Touchon M, Tourdjman M, Walker M, Stroika S, Cantinelli T, Chenal-Francisque V, Kucerova Z, Rocha EPC, Nadon C, Grant K, Nielsen EM, Pot B, Gerner-Smidt P, Lecuit M, Brisse S. 2016. Whole genome-based population biology and epidemiological surveillance of *Listeria monocytogenes*. Nat Microbiol 2:1–10.

50. de Been M, Pinholt M, Top J, Bletz S, Mellmann A, van Schaik W, Brouwer E, Rogers M, Kraat Y, Bonten M, Corander J, Westh H, Harmsen D, Willems RJL. 2015. Core Genome Multilocus Sequence Typing Scheme for High-Resolution Typing of *Enterococcus faecium*. J Clin Microbiol 53:3788–3797.

51. Cody AJ, Bray JE, Jolley KA, McCarthy ND, Maiden MCJ. 2017. Core Genome Multilocus Sequence Typing Scheme for Stable, Comparative Analyses of Campylobacter jejuni and C. coli Human Disease Isolates. J Clin Microbiol 55:2086–2097.

52. Janowicz A, De Massis F, Ancora M, Camma C, Patavino C, Battisti A, Prior K, Harmsen D, Scholz H, Zilli K, Sacchini L, Di Giannatale E, Garofolo G. 2018. Core Genome Multilocus Sequence Typing and Single Nucleotide Polymorphism Analysis in the Epidemiology of *Brucella melitensis* Infections. J Clin Microbiol 56:e00517–18.

53. Bletz S, Janezic S, Harmsen D, Rupnik M, Mellmann A. 2018. Defining and Evaluating a Core Genome Multilocus Sequence Typing Scheme for Genome-Wide Typing of *Clostridium difficile*. J Clin Microbiol 56:e01987–17.

54. Jones RC, Harris LG, Morgan S, Ruddy MC, Perry M, Williams R, Humphrey T, Temple M, Davies AP. 2019. Phylogenetic Analysis of *Mycobacterium tuberculosis* Strains in Wales by Use of Core Genome Multilocus Sequence Typing To Analyze Whole-Genome Sequencing Data. J Clin Microbiol 57:e02025–18.

55. Sails AD, Swaminathan B, Fields PI. 2003. Clonal complexes of *Campylobacter jejuni* identified by multilocus sequence typing correlate with strain associations identified by multilocus enzyme electrophoresis. J Clin Microbiol 41:4058–4067.

56. Leavis HL, Bonten MJ, Willems RJ. 2006. Identification of high-risk enterococcal clonal complexes: global dispersion and antibiotic resistance. Curr Opin Microbiol 9:454–460.

57. Weill F-X, Domman D, Njamkepo E, Tarr C, Rauzier J, Fawal N, Keddy KH, Salje H, Moore S, Mukhopadhyay AK, Bercion R, Luquero FJ, Ngandjio A, Dosso M, Monakhova E, Garin B, Bouchier C, Pazzani C, Mutreja A, Grunow R, Sidikou F, Bonte L, Breurec S, Damian M, Njanpop-Lafourcade B-M, Sapriel G, Page A-L, Hamze M, Henkens M, Chowdhury G, Mengel M, Koeck J-L, Fournier J-M, Dougan G, Grimont PAD, Parkhill J, Holt KE, Piarroux R, Ramamurthy T, Quilici M-L, Thomson NR. 2017. Genomic history of the seventh pandemic of cholera in Africa. Science 358:785–789.

58. Reimer A, Domselaar G, Stroika S, Walker M, Kent H, Tarr C, Talkington D, Rowe L, Olsen-Rasmussen M, Frace M, Sammons S, Dahourou G, Boncy J, Smith A, Mabon P, Petkau A, Graham M, Gilmour M, Gerner-Smidt P. 2011. Comparative Genomics of *Vibrio cholerae* from Haiti, Asia, and Africa. Emerg Infect Dis 17:2113.

59. Salim A, Lan R, Reeves PR. 2005. *Vibrio cholerae* pathogenic clones. Emerg Infect Dis 11:1758–1760.

60. Dunn JC. 1974. Well-separated clusters and optimal fuzzy partitions. J Cybern 4:95–104.

61. Mutreja A, Kim DW, Thomson NR, Connor TR, Lee JH, Kariuki S, Croucher NJ, Choi SY, Harris SR, Lebens M, Niyogi SK, Kim EJ, Ramamurthy T, Chun J, Wood JLN, Clemens JD, Czerkinsky C, Nair GB, Holmgren J, Parkhill J, Dougan G. 2011. Evidence for several waves of global transmission in the seventh cholera pandemic. Nature 477:462–465.

62. Hubert L, Arabie P. 1985. Comparing partitions. J Classif 2:193–218.

63. Lucidarme J, Hill DMC, Bratcher HB, Gray SJ, du Plessis M, Tsang RSW, Vazquez JA, Taha M-K, Ceyhan M, Efron AM, Gorla MC, Findlow J, Jolley KA, Maiden MCJ, Borrow R. 2015. Genomic resolution of an aggressive, widespread, diverse and expanding meningococcal serogroup B, C and W lineage. J Infect 71:544–552.

64. Royer G, Fourreau F, Boulanger B, Mercier-Darty M, Ducellier D, Cizeau F, Potron A, Podglajen I, Mongardon N, Decousser J-W. 2019. Local outbreak of extended-spectrum β-lactamase SHV2a-producing *Pseudomonas aeruginosa* reveals the emergence of a new specific sub-lineage of the international ST235 high-risk clone. J Hosp Infect.

65. Eppinger M, Pearson T, Koenig SSK, Pearson O, Hicks N, Agrawal S, Sanjar F, Galens K, Daugherty S, Crabtree J, Hendriksen RS, Price LB, Upadhyay BP, Shakya G, Fraser CM, Ravel J, Keim PS. 2014. Genomic Epidemiology of the Haitian Cholera Outbreak: a Single Introduction Followed by Rapid, Extensive, and Continued Spread Characterized the Onset of the Epidemic. MBio 5:e01721–14.

66. World Health Organization. 2017. Weekly epidemiological record.

67. Cavailler P, Lucas M, Perroud V, McChesney M, Ampuero S, Guérin PJ, Legros D, Nierle T, Mahoudeau C, Lab B, Kahozi P, Deen JL, von Seidlein L, Wang XY, Puri M, Ali M, Clemens JD, Songane F, Baptista A, Ismael F, Barreto A, Chaignat CL. 2006. Feasibility of a mass vaccination campaign using a two-dose oral cholera vaccine in an urban cholera-endemic setting in Mozambique. Vaccine 24:4890–4895.

68. Sardar T, Mukhopadhyay S, Bhowmick AR, Chattopadhyay J. 2013. An optimal cost effectiveness study on Zimbabwe cholera seasonal data from 2008-2011. PLoS One 8:e81231.

69. Hasan NA, Choi SY, Eppinger M, Clark PW, Chen A, Alam M, Haley BJ, Taviani E, Hine E, Su Q, others. 2012. Genomic diversity of 2010 Haitian cholera outbreak strains. Proc Natl Acad Sci 109:E2010–E2017.

70. Hu D, Liu B, Feng L, Ding P, Guo X, Wang M, Cao B, Reeves PR, Wang L. 2016. Origins of the current seventh cholera pandemic. Proc Natl Acad Sci 113:E7730–E7739.

71. Guillaume Y, Ternier R, Vissieres K, Casseus A, Chery MJ, Ivers LC. 2018. Responding to cholera in Haiti: Implications for the national plan to eliminate cholera by 2022. J Infect Dis 218:S167–S170.

72. Pightling AW, Petronella N, Pagotto F. 2014. Choice of Reference Sequence and Assembler for Alignment of *Listeria monocytogenes* Short-Read Sequence Data Greatly Influences Rates of Error in SNP Analyses. PLoS One 9:e104579.

73. Childers BM, Klose KE. 2007. Regulation of virulence in Vibrio cholerae: The ToxR regulon. Future Microbiol 2:335–344.

74. Souvorov A, Agarwala R, Lipman DJ. 2018. SKESA: strategic k-mer extension for scrupulous assemblies. Genome Biol 19:153.

75. Edgar RC. 2010. Search and clustering orders of magnitude faster than BLAST. Bioinformatics 26:2460–2461.

76. Aziz RK, Bartels D, Best AA, DeJongh M, Disz T, Edwards RA, Formsma K, Gerdes S, Glass EM, Kubal M, Meyer F, Olsen GJ, Olson R, Osterman AL, Overbeek RA, McNeil LK, Paarmann D, Paczian T, Parrello B, Pusch GD, Reich C, Stevens R, Vassieva O, Vonstein V, Wilke A, Zagnitko O, Formsma K, Kubal M, Vonstein V, Stevens R, McNeil LK, Edwards RA, Pusch GD, Reich C, Glass EM, Olsen GJ, Paczian T, Overbeek RA, Meyer F, Vassieva O, DeJongh M, Osterman AL, Disz T, Best AA, Gerdes S, Parrello B, Bartels D, Olson R, Paarmann D. 2008. The RAST server: rapid annotations using subsystems technology. BMC Genomics 9:75.

77. Islam MT, Liang K, Im MS, Winkjer J, Busby S, Tarr CL, Boucher Y. 2018. Draft Genome Sequences of Nine *Vibrio* sp. Isolates from across the United States Closely Related to Vibrio cholerae. Microbiol Resour Announc 7:e00965–18.

78. Liang K, Islam MT, Hussain N, Winkjer NS, Im MS, Rowe LA, Tarr CL, Boucher Y. 2019. Draft Genome Sequences of Eight Vibrio sp. Clinical Isolates from across the United States That Form a Basal Sister Clade to *Vibrio cholerae*. Microbiol Resour Announc 8:e01473–18.

79. Liang K, Orata FD, Winkjer NS, Rowe LA, Tarr CL, Boucher Y. 2017. Complete Genome Sequence of *Vibrio* sp. Strain 2521-89, a Close Relative of *Vibrio cholerae* Isolated from Lake Water in New Mexico, USA. Genome Announc 5:e00905–17.

80. Goris J, Konstantinidis KT, Klappenbach JA, Coenye T, Vandamme P, Tiedje JM. 2007. DNA–DNA hybridization values and their relationship to whole-genome sequence similarities. Int J Syst Evol Microbiol 57:81–91.

81. Meier-Kolthoff JP, Auch AF, Klenk HP, Göker M. 2013. Genome sequence-based species delimitation with confidence intervals and improved distance functions. BMC Bioinformatics 14:60.

82. Parks DH, Imelfort M, Skennerton CT, Hugenholtz P, Tyson GW. 2015. CheckM : assessing the quality of microbial genomes recovered from isolates, single cells, and metagenomes. Genome Res 25:1043–1055.

83. Altschul SF, Gish W, Miller W, Myers EW, Lipman DJ. 1990. Basic local alignment search tool. J Mol Biol 215:403–410.

84. Jolley KA, Maiden MCJ. 2010. BIGSdb: Scalable analysis of bacterial genome variation at the population level. BMC Bioinformatics 11:595.

85. Garg P, Aydanian A, Smith D, Morris JGJG, Nair GB, Stine OC, Antonia A, Smith D, Morris JGJG, Nair GB, Stine OC. 2003. Molecular Epidemiology of O139 *Vibrio cholerae*: Mutation, Lateral Gene Transfer, and Founder Flush. Emerg Infect Dis 9:810–814.

86. Brock G, Pihur V, Datta S, Datta S. 2008. clValid: An R Package for Cluster Validation. J Stat Softw 25:1–22.

87. R Core Team. 2017. R: A Language and Environment for Statistical Computing. Vienna, Austria.

88. Canty A, Ripley BD, others. 2017. boot: Bootstrap R (S-Plus) Functions. R Packag version 1.

89. Wickham H. 2009. ggplot2: Elegant Graphics for Data Analysis. Springer-Verlag New York.

90. Chang F, Qiu W, Zamar RH, Lazarus R, Wang X. 2010. clues: An R Package for Nonparametric Clustering Based on Local Shrinking. J Stat Softw 33:1–16.

91. Zhou Z, Alikhan N, Sergeant MJ, Luhmann N, Vaz C, Francisco AP, Carriço JA, Achtman M. 2018. GrapeTree: visualization of core genomic relationships among 100,000 bacterial pathogens. Genome Res 28:1395–1404.

92. Treangen TJ, Ondov BD, Koren S, Phillippy AM. 2014. The harvest suite for rapid core-genome alignment and visualization of thousands of intraspecific microbial genomes. Genome Biol 15:524.

93. Bruen TC, Philippe H, Bryant D. 2006. A simple and robust statistical test for detecting the presence of recombination. Genetics 172:2665–2681.

94. Letunic I, Bork P. 2007. Interactive Tree Of Life (iTOL): An online tool for phylogenetic tree display and annotation. Bioinformatics 23:127–128.

95. Hagberg A, Swart P, S Chult D. 2008. Exploring network structure, dynamics, and function using NetworkX, p. 11–16. *In* Varoquaux G, Vaught T, Millman J (ed), Proceedings of the 7th Python in Science Conference (SciPy2008).

96. Shannon P, Markiel A, Ozier O, Baliga NS, Wang JT, Ramage D, Amin N, Schwikowski B, Ideker T. 2003. Cytoscape: A Software Environment for Integrated Models of Biomolecular Interaction Networks. Genome Res 13:2498–2504.

