## Supplemental Figure S1-S3 for "A *Vibrio cholerae* Core Genome Multilocus Sequence Typing Scheme to Facilitate the Epidemiological Study of Cholera"

### SUPPLEMENTAL MATERIAL

Kevin Y. H. Liang,[a] Fabini D. Orata,[a] Mohammad Tarequl Islam,[a] Tania Nasreen,[a] Munirul Alam,[b] Cheryl L. Tarr,[c] Yann F. Boucher[a#]

[a]Department of Biological Sciences, University of Alberta, Edmonton, Alberta, Canada
[b]Infectious Diseases Division, International Centre for Diarrhoeal Disease Research, Dhaka, Bangladesh
[c]Enteric Diseases Laboratory Branch, Centers for Disease Control and Prevention, Atlanta, Georgia, USA

**Supplemental Figures**

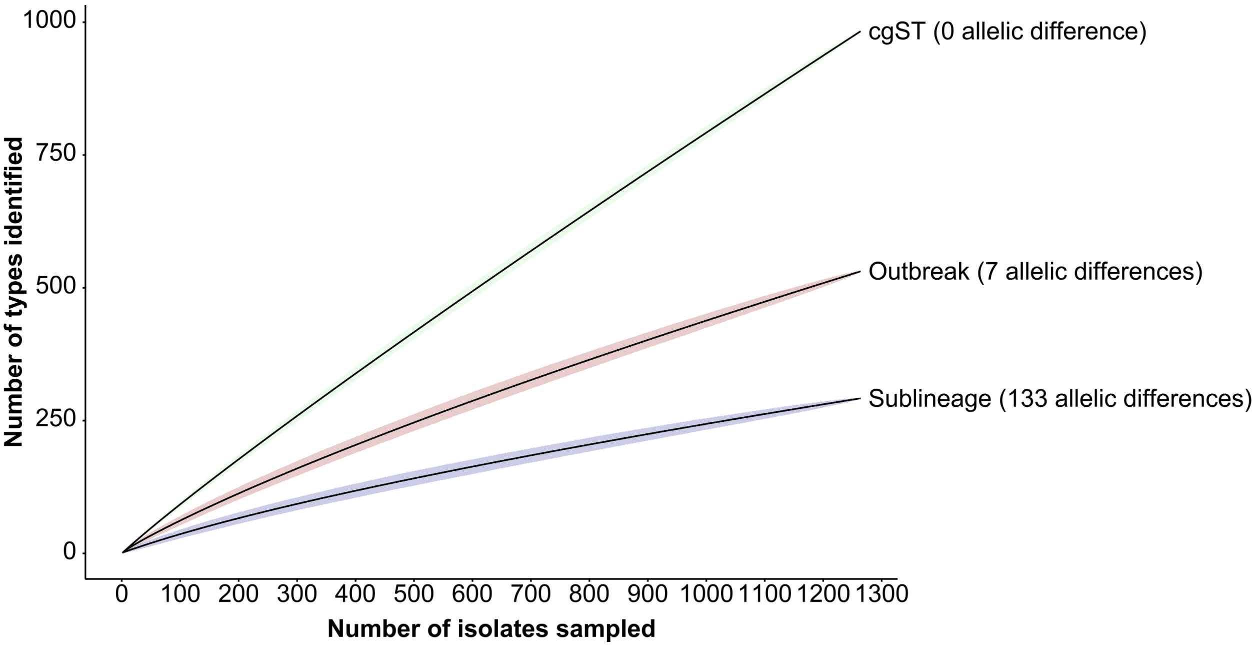


**Fig. S1.** Rarefaction curve for cgST, outbreak threshold (7 allelic differences), and the sublineage threshold (133 allelic differences) computed using mothur (1) with default parameters.

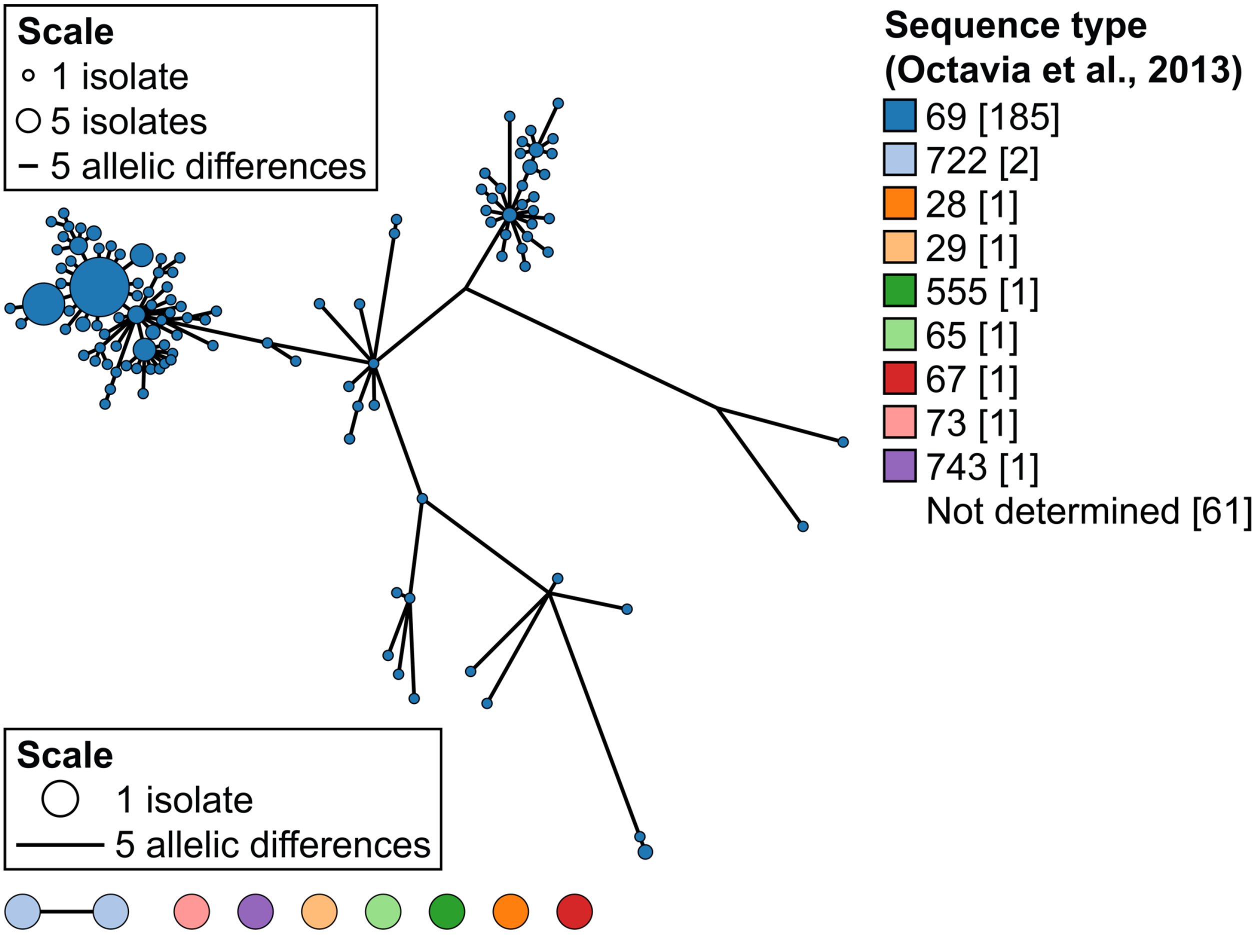


**Fig. S2.** Evaluation of network similarities between the cgMLST sublineage threshold and MLST ST. Networks of all sublineages identified using only *V. cholerae* isolates from Bangladesh ($n$ = 255). Each cluster represents a sublineage and includes isolates with less than or equal to 133 allelic differences with each other. Each node represents a cgST and is colored by sequence type based on the 2013 MLST scheme (2). Size of the nodes are proportional to the number of isolates. The length of the connecting lines within a cluster is proportional to the number of allelic differences.

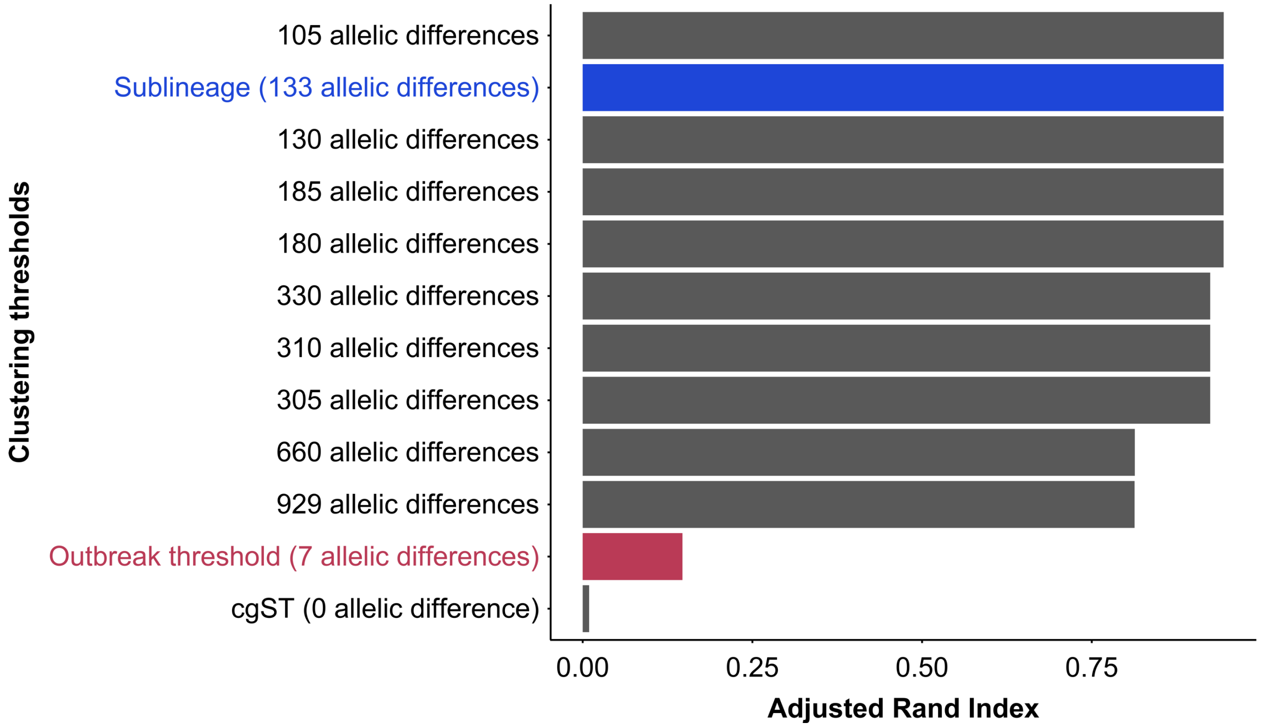


**Fig. S3.** Adjusted Rand Index for individual pairwise comparisons between predefined clustering thresholds (Fig. 2) and the 2013 MLST scheme (2). The sublineage clustering threshold (i.e., 133 allelic differences) and outbreak threshold (i.e., 7 allelic differences) are indicated in blue and red bars, respectively.

**Supplemental Tables***

**Table S1.** Meta-information for all 1,262 isolates used in this study.

**Table S2.** Allelic profiles for the cgMLST scheme (as defined in this study), the 2016 MLST scheme by Kirchberger and colleagues (3), and the 2013 MLST scheme by Octavia and colleagues (2). The cgSTs and their corresponding STs and PubMLST IDs are indicated. All missing genes are indicated as NA. The most likely cgSTs are indicated in parenthesis where applicable.

**Table S3.** *V. cholerae* isolates from the Yemen cholera outbreak and neighbouring countries, as well as other isolates from different lineages.

**Table S4.** Genome completeness for the cgMLST scheme (using 2,443 core genes). All genomes with less than 90% completeness were subsequently removed.

**Table S5.** Genome completeness information for the final set of 679 genomes. Completeness for the cgMLST scheme is represented as the percentage of the of the 2,443 core genes present in each genome.

**Table S6.** All NCBI accession numbers for isolates, PubMLST IDs, and links to online storage of genomes.

**Please see separate Excel sheets for complete supplemental tables.*

**References**

1. Schloss PD, Westcott SL, Ryabin T, Hall JR, Hartmann M, Hollister EB, Lesniewski RA, Oakley BB, Parks DH, Robinson CJ, Sahl JW, Stres B, Thallinger GG, Van Horn DJ, Weber CF. 2009. Introducing mothur: open-source, platform-independent, community-supported software for describing and comparing microbial communities. Appl Environ Microbiol 75:7537–41.

2. Octavia S, Salim A, Kurniawan J, Lam C, Leung Q, Ahsan S, Reeves PR, Nair GB, Lan R. 2013. Population structure and evolution of non-O1/non-O139 *Vibrio cholerae* by multilocus sequence typing. PLoS One 8:e65342.

3. Kirchberger PC, Orata FD, Barlow EJ, Kauffman KM, Case RJ, Polz MF, Boucher Y. 2016. A small number of phylogenetically distinct clonal complexes dominate a coastal *Vibrio cholerae* population. Appl Environ Microbiol 82:5576–5586.
